# Evaluating the added value of sequence models in noncoding RNA assays: a baseline-aware framework

**DOI:** 10.64898/2026.10.22.760090

**Authors:** Rui Guo, Wenhui Shan, Ruoyan Liu, Lei Yang, Dongmei Mu, Hong Wang

## Abstract

Sequence models can improve prediction of non-coding RNA assay read-outs, but their contribution depends on the comparison baseline. We present a baseline-aware framework that estimates model-added-last changes in AUROC for lncRNA labels and cross-validated Spearman correlation for miRNA repression, fixing members and chromosome folds across available sources. In cross-cell CRISPRi (n = 2961), AlphaGenome increased AUROC from 0.6426 to 0.7551; adding K562 accessibility raised baseline AUROC to 0.7282, with augmented AUROC of 0.7534. Corrected-source Cas13 increments ranged from –0.0161 to +0.0130 (n = 871). Post-audit comparisons evaluated sequence composition, broader regularization and TargetScan aggregate identity. With expanded regularization, composition improved four baselines but reduced Cas13 AUROC by 0.0365. After composition adjustment, RNA-FM’s increment was –0.0014 for CRISPRi under the initial grid and +0.000044 for HEK293FT under the expanded grid. Small increments changed sign with regularization. On the same 1227 HeLa members, replacing a seed-family aggregate with an identity-supported mature-miRNA aggregate increased baseline Spearman from 0.4140 to 0.4799. RNA-FM added 0.0170 beyond this baseline, or 0.0014 after composition adjustment. These gene-level aggregates lacked experimental transcript/site alignment. Resampling was descriptive or non-joint, without calibrated confirmation, prespecified practical-size thresholds or verified pretraining overlap. The framework identifies positive model contributions and makes their dependence on baseline content, feature identity and regularization explicit, providing a practical basis for interpreting sequence-model performance.

## Background

Deep-learning sequence models predict genomic and transcriptomic measurements. AlphaGenome, Enformer and Borzoi map genomic windows to regulatory tracks including chromatin accessibility and transcription and, for Borzoi, RNA-seq coverage [1–3]; genomic language models remain under active development [4]. RNA-FM learns transcript representations from unannotated RNA collections [5] and has been benchmarked on RNA structure and function tasks [6]. Such representations are also tested against non-coding RNA (ncRNA) assays. Here, the labels are CRISPRi lncRNA growth or fitness [7] and quantitative miRNA repression with TargetScan features [8, 9]. Related benchmarks address RNA processes [10], lncRNA coding potential [11] and 3’UTR prediction [12]. These benchmarks compare performance on specific tasks. We ask how much a named model feature block adds over an explicit baseline on the same assay members. We then examine how that estimate changes with the baseline.

Explicit biological baselines help interpret what sequence models contribute to assay prediction. Sequence models more readily recover promoter-proximal expression determinants than distal effects [13] and explain limited between-individual expression variation, including cis-regulatory variant directions [14, 15]. Transcript abundance and accessibility may correlate with lncRNA growth or fitness, while target-site context may correlate with miRNA repression. We therefore distinguish minimally adjusted from accessibility-adjusted estimands. Accessibility may act as a proxy, mediator or both. These adjustment sets therefore define complementary predictive questions; partitioning cis and trans components would require additional identification assumptions.

Reliable downstream evaluation keeps supervised probes, reduction and tuned transformations within outer-training folds [16]. This protects the outer test from adaptive fitting; overlap with foundation-model pretraining is a separate provenance question. Classification and continuous outcomes also require different metrics; a common display scale would obscure this distinction. We retain native delta AUROC and delta cross-validated Spearman, report adjustment sets, and use negative controls to examine specified failure modes [17].

The protocol follows added-predictive-value methods [18, 19] with outer-fold evaluation and internal grouped fitting. Each member fixes its label, nuisance covariates, folds and row identifiers before adding a model block. LncRNA cells use locus-level resampling and a locus-level null. MiRNA cells use separate transcript– and guide-cluster arms. Their interval envelope and diagnostic tail-area maximum describe sensitivity to each clustering dimension separately; crossed covariance is not estimated. Equivalence requires a prospectively justified, task-native smallest effect size of interest [20,21]; this study reports predictive increments, with practical-size assessment reserved for a design that specifies this threshold.

Worked examples cover lncRNA CRISPRi and Cas13 growth or fitness and miRNA repression, using named score sources where available. Together, the workflow (Box 1; Fig. 1a), results overview (Table 1) and decision aid (Box 2; Fig. 1b) show how to quantify model contributions and examine their sensitivity to the biological baseline. The comparisons concern the evaluated sources and assays, with model-class rankings and mechanistic questions requiring broader designs.

**Figure 1.**
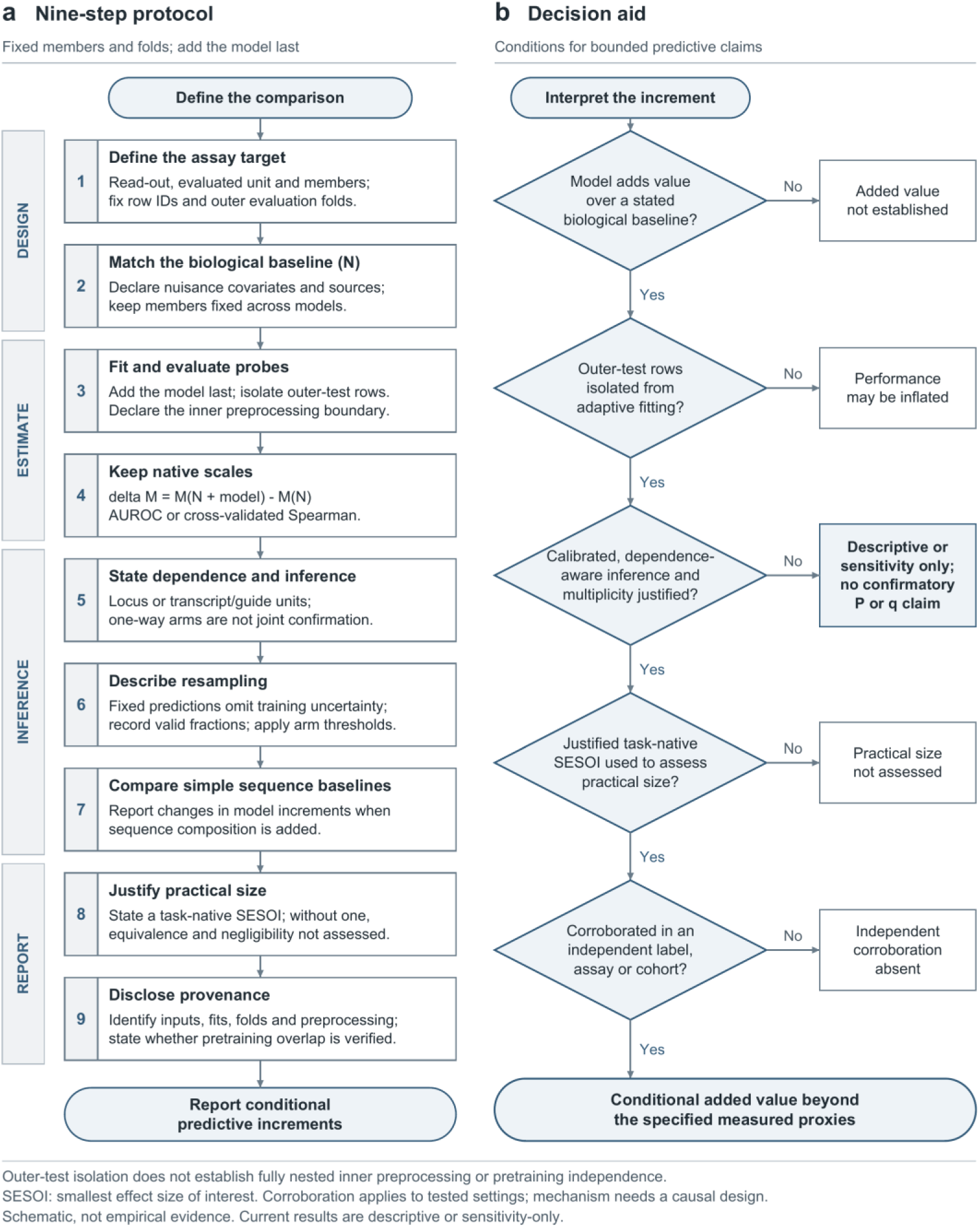
Workflow and interpretation of conditional predictive increments. The schematic summarizes the evaluation design. (a) The nine-step workflow records the assay, biological baseline, cross-fitted probe and inner preprocessing boundary, native increment, dependence and inference status, fixed-prediction resampling role, simple-sequence baseline sensitivity, practical-size status and provenance. (b) The decision aid first asks whether model features are compared with an explicit baseline, then checks outer-test isolation and declared inner preprocessing, justified inference, a smallest effect of interest, and independent corroboration. Outer-test isolation, inner preprocessing and pretraining overlap are separate checks. The present analysis follows the descriptive and sensitivity reporting branches.

**Table 1.** Absolute baseline and model augmented performance.

| Member | Model | n | Baseline | Augmented | Native increment | Range or one way envelope |
| --- | --- | --- | --- | --- | --- | --- |
| Cas13 | AG | 871 | 0.5898 | 0.6028 | +0.0130 | -0.0248 to +0.0475 |
| Cas13 | Borzoi | 871 | 0.5898 | 0.5787 | -0.0111 | -0.0258 to +0.0035 |
| Cas13 | RNA-FM | 871 | 0.5898 | 0.5778 | -0.0120 | -0.0235 to -0.0006 |
| Cas13 | Enformer | 871 | 0.5898 | 0.5738 | -0.0161 | -0.0293 to -0.0042 |
| CRISPRi minimal | AG | 2961 | 0.6426 | 0.7551 | +0.1125 | +0.0680 to +0.1537 |
| CRISPRi minimal | Borzoi | 2961 | 0.6426 | 0.7378 | +0.0953 | +0.0629 to +0.1293 |
| CRISPRi minimal | Enformer | 2961 | 0.6426 | 0.7255 | +0.0830 | +0.0503 to +0.1153 |
| CRISPRi minimal | RNA-FM | 2961 | 0.6426 | 0.6877 | +0.0452 | +0.0137 to +0.0780 |
| CRISPRi accessibility | AG | 2961 | 0.7282 | 0.7534 | +0.0252 | +0.0094 to +0.0412 |
| CRISPRi accessibility | Borzoi | 2961 | 0.7282 | 0.7528 | +0.0246 | +0.0101 to +0.0382 |
| CRISPRi accessibility | RNA-FM | 2961 | 0.7282 | 0.7467 | +0.0184 | +0.0040 to +0.0328 |
| CRISPRi accessibility | Enformer | 2961 | 0.7282 | 0.7454 | +0.0171 | +0.0055 to +0.0282 |
| HeLa disruption | AG | 2944 | 0.4201 | 0.4208 | +0.0008 | -0.0005 to +0.0020 |
| HeLa expression | AG | 2944 | 0.4201 | 0.4197 | -0.0004 | -0.0013 to +0.0004 |
| HEK293FT | RNA-FM | 2726 | 0.3421 | 0.3635 | +0.0214 | +0.0107 to +0.0306 |
| HEK293FT disruption | AG | 2726 | 0.3421 | 0.3454 | +0.0034 | -0.0013 to +0.0076 |
| HEK293FT expression | AG | 2726 | 0.3421 | 0.3413 | -0.0008 | -0.0022 to +0.0010 |
| HeLa | RNA-FM | 2944 | 0.4201 | 0.4203 | +0.0002 | -0.0012 to +0.0015 |

These 18 entries reproduce R6 and pair each increment with its absolute baseline and augmented performance. LncRNA uses AUROC and miRNA uses Spearman on separate scales. Ranges condition on fixed predictions and exclude training uncertainty; miRNA envelopes summarize separate one-way sensitivities. The CRISPRi outcome pools strict hits across seven cell lines; accessibility denotes K562 covariates. Confirmatory inference is not assessed. Differences were calculated from unrounded estimates; subtraction of the displayed values may differ in the final digit because each value is rounded independently.

### Protocol

The common protocol element is an added-last contrast (Box 1), retained on each task’s native scale. Within each lncRNA member, label, nuisance covariates, folds and row identifiers were fixed across available model sources. For each miRNA cohort, we first took the common site-ID intersection of the AlphaGenome disruption track, AlphaGenome expression track and RNA-FM feature source. The outcome, nuisance covariates, folds, transcript clusters and guide clusters were then fixed on that shared footprint:

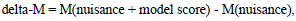

where M is AUROC for binary lncRNA labels and cross-validated Spearman correlation for continuous miRNA labels. Model transformations and fitting used outer-training rows only. The primary nuisance model was linear after outer-fold preprocessing; a polynomial model was a sensitivity. LncRNA summaries followed the locus comparator contract; miRNA summaries examined transcript– and guide-cluster sensitivity separately. The analysis targets native predictive increments. Practical equivalence would additionally require a locked task-native SESOI.

The benchmark comprised cross-cell CRISPRi strict lncRNA-hit labels with minimal covariates (n = 2961), the same labels with added K562 accessibility covariates, corrected-source Cas13 growth-or-fitness labels (n = 871), and common-footprint canonical-site miRNA repression in HeLa (n = 2944) and HEK293FT (n = 2726). The incomplete model-by-member matrix contains 18 declared comparisons (Table S1; Fig. 2). Absolute baseline and augmented performance are reported with each increment (Table 1; Table S2). Supplementary audit tables retain historical tail areas and BY/BH arithmetic with their non-confirmatory roles; unavailable miRNA inference is labelled not assessed. Synthetic spike-ins and negative controls check selected implementation paths (Table S3), complementing the observed-data comparisons.

**Figure 2.**
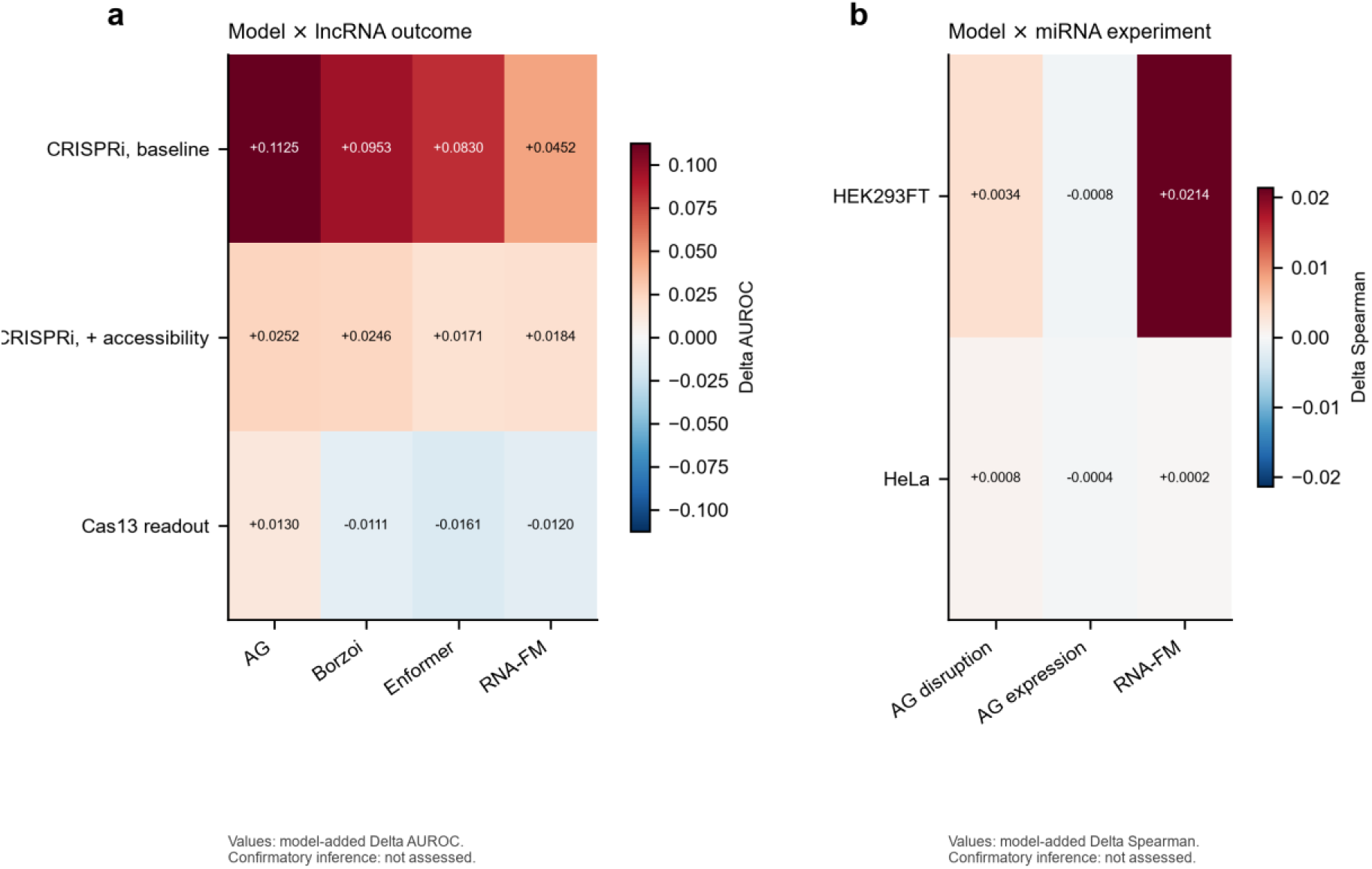
Benchmark landscape separated by native effect scale. (a) Heatmap of delta AUROC for binary lncRNA growth-or-fitness members. (b) Heatmap of delta cross-validated Spearman for continuous miRNA repression cohorts, with separate AlphaGenome disruption, AlphaGenome expression and RNA-FM columns. The panels display the evaluated model-by-member combinations on separate native metrics. Precision-recall sensitivities are reported separately, and the plotted contrasts address prediction rather than causal mediation.

### Case study 1 LncRNA growth or fitness readouts

For minimally adjusted cross-cell CRISPRi strict-hit labels, AlphaGenome increased AUROC from 0.6426 to 0.7551, with delta AUROC +0.1125 (descriptive fixed-prediction locus-resampling range +0.0680 to +0.1537). Estimates were +0.0953 for Borzoi, +0.0830 for Enformer and +0.0452 for RNA-FM (Table S1; Fig. 3a). All evaluated feature blocks contributed positive point increments on this baseline. The estimates describe the named sources in this assay; supplementary tail areas and BY/BH transformations retain their historical diagnostic roles.

**Figure 3.**
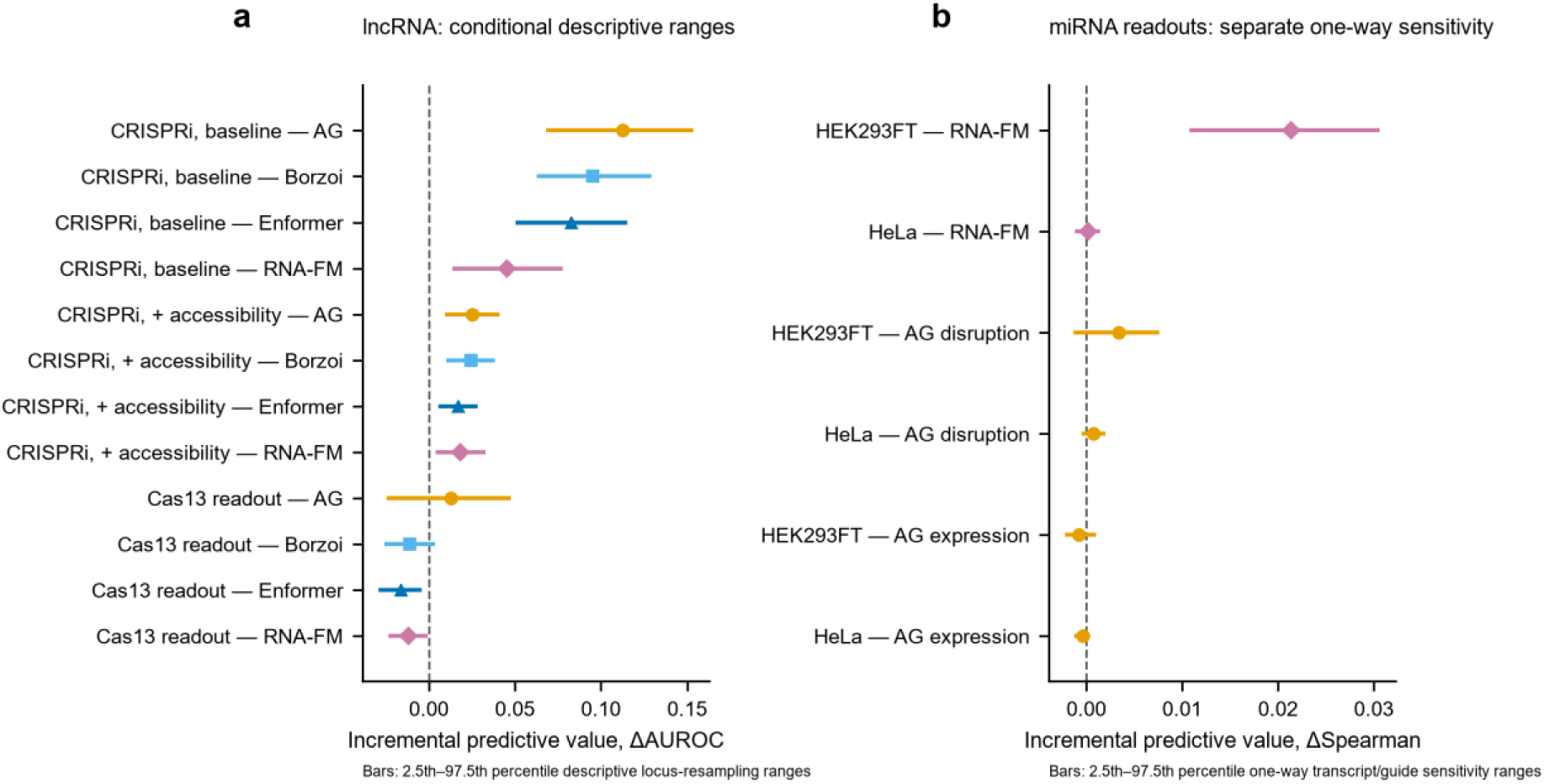
Native-scale added predictive value across assay members. Points show model-added-last estimates. (a) For the minimally adjusted, accessibility-adjusted and corrected Cas13 lncRNA growth-or-fitness members, horizontal lines describe locus resampling conditional on the realised fixed predictions. (b) For miRNA repression, the six common-footprint cells comprise RNA-FM and separate AlphaGenome disruption and expression tracks in HEK293FT and HeLa; horizontal lines are transcript/guide one-way sensitivity envelopes. Both range types are descriptive rather than confirmatory confidence intervals. Each panel uses its own native scale. Equivalence bands require a locked task-native SESOI and are omitted here.

Adding K562 promoter DNase and ATAC to the baseline changed the AlphaGenome estimate from +0.1125 to +0.0252. Baseline and augmented AUROC were 0.7282 and 0.7534 (descriptive 95% locus-resampling range conditional on fixed cross-fitted predictions, +0.0094 to +0.0412). Accessibility-adjusted Borzoi and Enformer estimates were +0.0246 and +0.0171 (Table S1). Positive increments remained after accessibility adjustment, while the higher baseline accounted for much of the difference in incremental performance. These analyses address distinct conditional questions: the labels pool cell lines and K562 accessibility supplies a single-context proxy. Separating mediation from confounding or localising cis and trans contributions requires a causal design. Fig. 2a displays the lncRNA native-scale comparisons. Precision-recall changes are secondary sensitivities reported separately from Fig. 2.

For the corrected Cas13 growth-or-fitness member (n = 871; 149 positives), the model-added-last delta AUROC estimates were +0.0130 for AlphaGenome, –0.0111 for Borzoi, – 0.0161 for the cached Enformer community port and –0.0120 for RNA-FM (Table S1). Their descriptive fixed-prediction locus-resampling ranges were –0.0248 to +0.0475, –0.0258 to +0.0035, –0.0293 to –0.0042 and –0.0235 to –0.0006, respectively. The ranges describe resampling of fixed predictions. The official TensorFlow Hub Enformer sensitivity was – 0.0037 (descriptive range –0.0319 to +0.0257; Table S4c). The official implementation supplies a provenance sensitivity within the same assay. Cas13 provides a complementary perturbation read-out to CRISPRi, while accessibility– and expression-related processes may still contribute to its labels. Interpretation remains descriptive, without a locked task-native SESOI. A later source-coverage sensitivity used retained Enformer and Borzoi community-port features on 946 genes/167 positives, adding 75 genes/18 positives to the shared footprint (Table S5). This extends coverage within E051. Under the expanded-grid fitting protocol, N and N+S AUROC were 0.5784 and 0.5481. Enformer increments were –0.0109/-0.0036 over N/N+S; Borzoi increments were –0.0019/-0.0002. The coverage comparison captures changes in both evaluation membership and the training population.

### Case study 2 MiRNA repression readouts

For the common-footprint HEK293FT canonical-site repression member (n = 2726), RNA-FM had delta cross-validated Spearman +0.0214; the one-way transcript/guide sensitivity envelope was +0.0107 to +0.0306 (Table S1). This positive point increment identifies an RNA-FM contribution under the declared HEK293FT baseline. The envelope describes separate clustering sensitivities; joint confirmation and practical-size assessment remain open because calibrated crossed inference and a locked SESOI were unavailable. The common-footprint HeLa estimate was +0.0002 (Fig. 3b), illustrating the assay dependence on the native Spearman scale.

The historical gene-by-seed-family TargetScan join covered 705 of 2726 HEK293FT rows and 1227 of 2944 HeLa rows. Let A, B and C denote RNA-FM’s added-last Spearman increment in the full cohort, in the covered subset with the original nuisance covariates, and in the same covered subset after adding the family aggregate, respectively. These increments were +0.0214, +0.0058 and +0.0012 in HEK293FT, and +0.0002, +0.0052 and –0.0029 in HeLa (Table S6a).

The covered-subset contrasts (B – A) were –0.0155 and +0.0050, and the within-subset adjustment contrasts (C – B) were –0.0047 and –0.0081, respectively (Table S6b). Contrasts were calculated from unrounded estimates and describe the contributions of subset selection and within-subset adjustment. The one-way resampling envelopes apply to individual increments, with no resampled inference for their differences. The family feature sums weighted context++ source rows by gene and seed family, potentially mixing mature miRNAs; its interpretation is therefore a family-level proxy for the experimental guide.

To assess the contribution of domain-feature identity, we refitted all three model branches on the same covered members with fixed alpha 100 (Table 2; Tables S7-S8; Fig. S1). Exact mature-guide matching supported 1227 HeLa and 645 HEK293FT rows. Replacing family aggregate F with identity-supported mature-miRNA aggregate G changed baseline Spearman from 0.4140 to 0.4799 in HeLa and from 0.4284 to 0.4358 in HEK293FT. With sequence composition S, the corresponding changes were 0.4417 to 0.4843 and 0.4386 to 0.4463. RNA-FM contributed positive increments of +0.0170/+0.0076 over N+G in HeLa/HEK293FT. With composition included, its increments were +0.0014/-0.0038 over N+S+G. The latter HEK293FT background retained AlphaGenome disruption/expression increments of +0.0051/+0.0024. These comparisons show how mature-guide identity strengthens the domain baseline and changes the residual contribution of each model block. Both F and G remain gene-level source-row sums without experimental transcript/site crosswalks. A separate 705-row HEK293FT sensitivity evaluates a terminal-U guide mapping with different membership; it addresses mapping and coverage together.

**Table 2.** Exact-identity TargetScan backgrounds and model increments (fixed alpha 100)

| Set | n | Background | Baseline rho | AG-disrupt<br>delta | AG-expression<br>delta | RNA-FM<br>delta |
| --- | --- | --- | --- | --- | --- | --- |
| hela_exact | 1227 | N__none | 0.3873 | +0.0003 | -0.0011 | +0.0306 |
| hela_exact | 1227 | N__family | 0.4140 | +0.0002 | -0.0009 | +0.0299 |
| hela_exact | 1227 | N__guide | 0.4799 | +0.000005 | -0.0005 | +0.0170 |
| hela_exact | 1227 | NS__none | 0.4199 | -0.0005 | +0.0008 | +0.0007 |
| hela_exact | 1227 | NS__family | 0.4417 | -0.0004 | +0.0009 | +0.0003 |
| hela_exact | 1227 | NS__guide | 0.4843 | -0.0003 | +0.0015 | +0.0014 |
| hek_exact | 645 | N__none | 0.3777 | +0.0056 | +0.0015 | +0.0212 |
| hek_exact | 645 | N__family | 0.4284 | +0.0051 | +0.0017 | +0.0108 |
| hek_exact | 645 | N__guide | 0.4358 | +0.0046 | +0.0025 | +0.0076 |
| hek_exact | 645 | NS__none | 0.3922 | +0.0064 | +0.0015 | -0.0038 |
| hek_exact | 645 | NS__family | 0.4386 | +0.0055 | +0.0022 | -0.0024 |
| hek_exact | 645 | NS__guide | 0.4463 | +0.0051 | +0.0024 | -0.0038 |

On the common footprint, the AlphaGenome disruption/expression delta cross-validated Spearman estimates were +0.0034/-0.0008 in HEK293FT and +0.0008/-0.0004 in HeLa (Table S1; Figs. 2b and 3b). The disruption track yielded positive point increments in both cohorts, alongside slightly negative expression-track estimates. These descriptive comparisons concern the evaluated AlphaGenome tracks and RNA-FM; Enformer and Borzoi were not evaluated in these members.

### Sensitivity analyses of the miRNA increment

Because chromosome folds can place rows from one transfected guide in training and test sets, a leave-one-guide-out sensitivity held out each guide and all its rows. The estimates were +0.0153 in HEK293FT and +0.0002 in HeLa (Table S9; Fig. 4a), with the same non-joint sensitivity role as primary miRNA rows. The positive HEK293FT point increment persisted when guides were held out, supporting its relevance beyond the primary chromosome partition. TargetScan mapped seed-family labels to 12 HEK293FT guides and 16 of 17 HeLa guides; lsy-6 remained the non-human control. Table S10 documents whether guide hold-out also separates every mapped seed family. This check addresses the tested partitions; broader family transfer, sequence similarity and pretraining overlap require separate evaluation.

**Figure 4.**
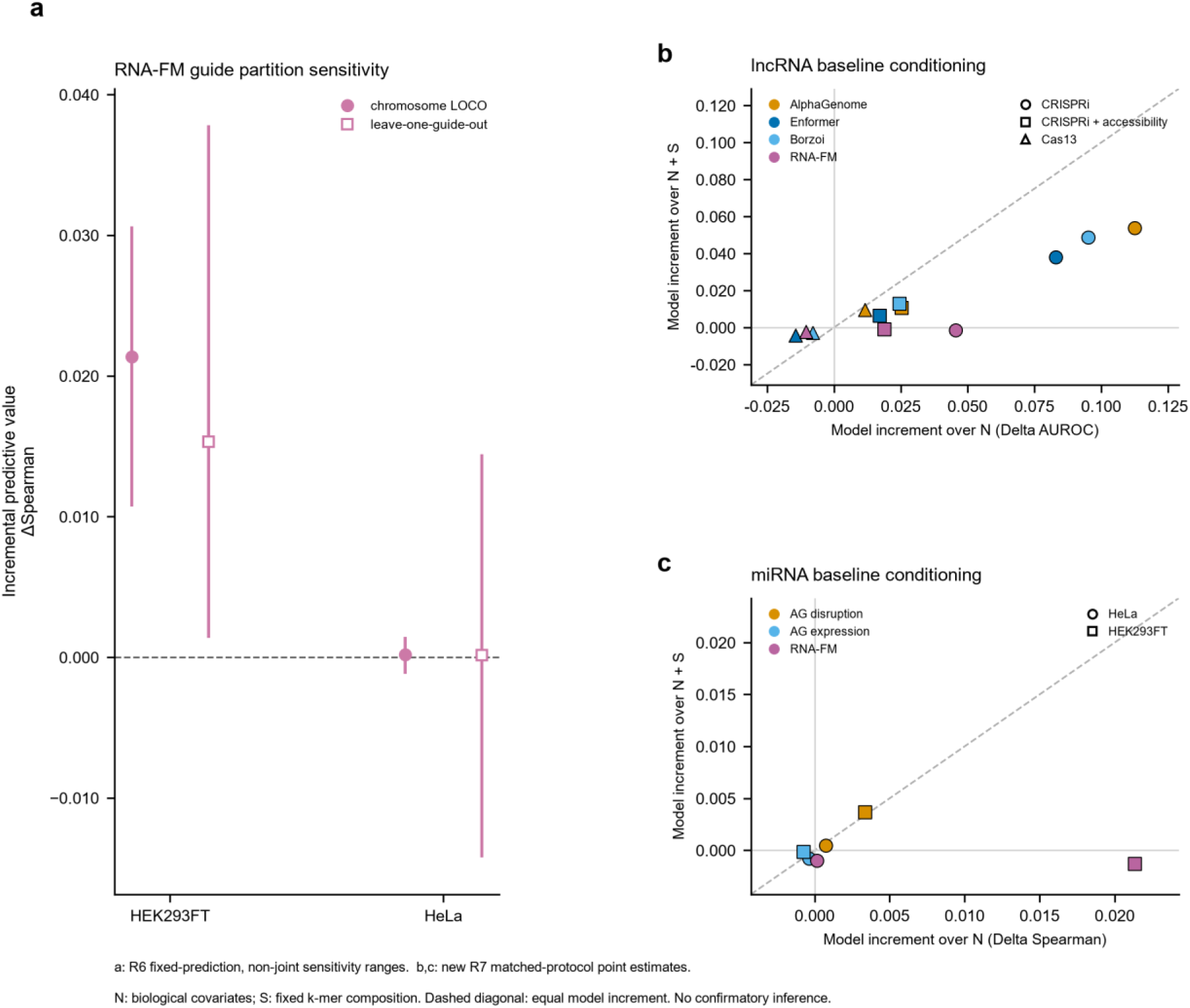
Guide partitioning and sequence-baseline sensitivity. (a) RNA-FM native delta cross-validated Spearman under primary chromosome folds and leave-one-guide-out folds in HEK293FT and HeLa, with descriptive transcript/guide one-way sensitivity envelopes. (b) Twelve lncRNA model cells and (c) six miRNA model cells compare the model increment given the measured baseline N on the horizontal axis with its increment given N plus sequence composition S on the vertical axis. These points use the matched new strict-inner fitting protocol; lncRNA AUROC and miRNA Spearman panels remain on separate native scales. The diagonal indicates equal point increments. The plots retain negative values and compare matched fits within the new composition protocol, separately from historical R6. Panels b,c retain the initial R7 restricted-grid estimates: all composition-baseline folds selected alpha 100. Tables S16 and S17 report the common-alpha and later expanded-grid diagnostics, respectively. Producer-PCA consistency is documented in Table S11. The envelopes and point comparisons are descriptive; practical equivalence is not assessed.

High-dimensional embeddings underwent outer-training-fold reduction before supervised fitting. Deterministic recomputation of outer-fold PCA matched the stored primary point estimates for every member. This verifies consistency of the recorded PCA point-estimation path (Table S11). At k = 3, 8, 10 and 16, the HEK293FT RNA-FM estimates were +0.0123, +0.0110, +0.0104, +0.0214; the corrected Cas13 estimates were –0.0129, +0.0040, –0.0038 and –0.0120 (Fig. 5a,b; Table S12). Representation sensitivities for corrected Cas13 were – 0.0120, +0.0060, +0.0093 and –0.0089 for full-transcript, 5-prime, centre and 3-prime summaries; HEK293FT target-only and target-plus-guide estimates were +0.0306 and +0.0214 (Fig. 5c,d; Table S13). Positive HEK293FT estimates across these downstream choices complement the primary result, while the Cas13 values show greater sensitivity to representation. Synthetic spike-ins and one-way guide diagnostics (Table S3), classification negative controls (Fig. 6a), and collinearity stress (Fig. 6c,d; Tables S3 and S14) examine specified implementation settings. Fig. 6b records the absence of registered miRNA negative-control rows. These diagnostics address implementation consistency; calibration and pretraining independence remain open questions, as described in the study limitations.

**Figure 5.**
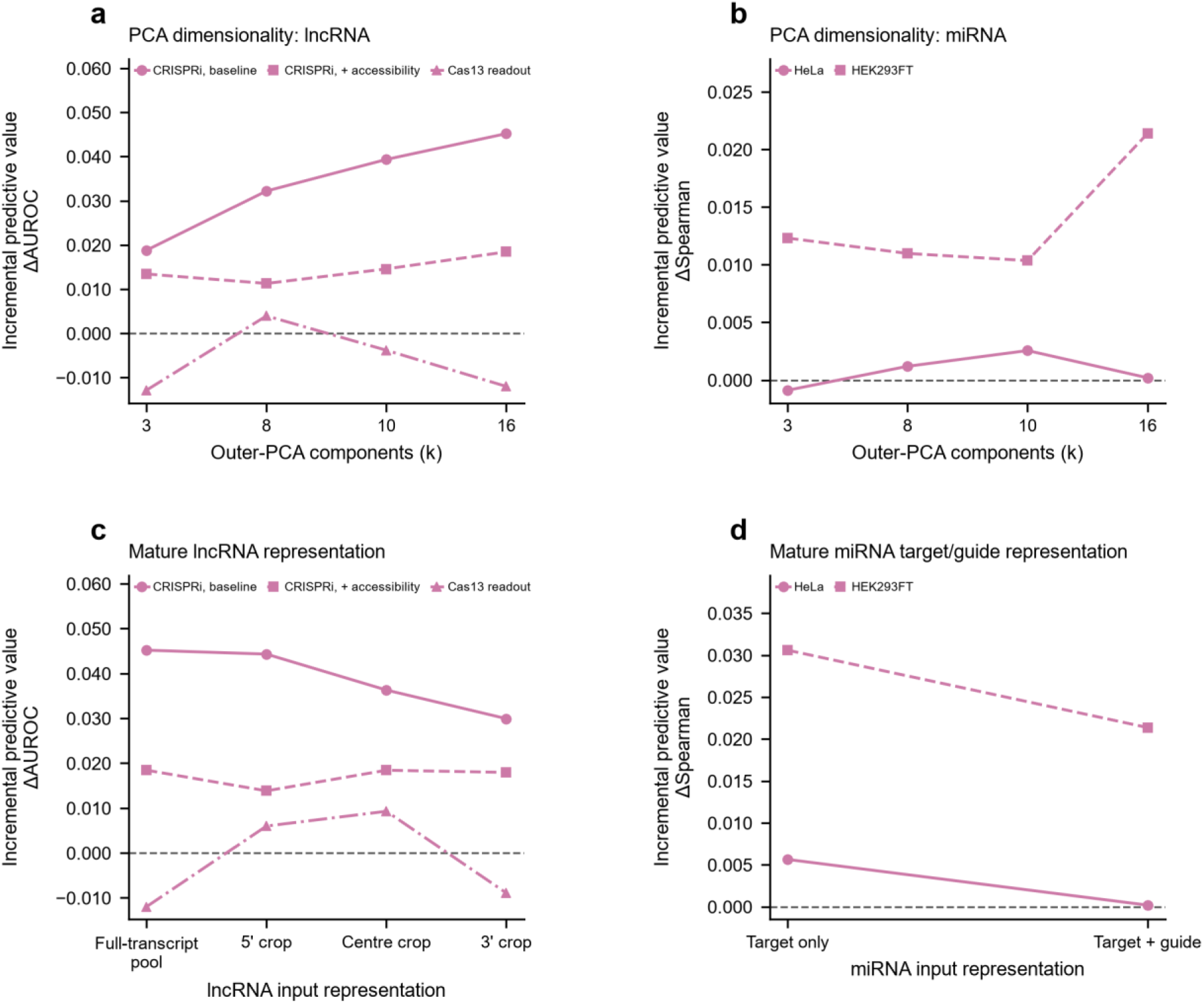
Downstream PCA and input-representation sensitivities. Native-scale RNA-FM point estimates across declared outer-fold PCA component counts are shown separately for (a) lncRNA members on delta AUROC and (b) miRNA cohorts on delta cross-validated Spearman. Input-representation point sensitivities compare (c) full-transcript nucleotide-weighted pooling with positional lncRNA summaries and (d) target-only with target-plus-guide paired miRNA features. The panels show how declared downstream choices affect point estimates around the primary analysis.

**Figure 6.**
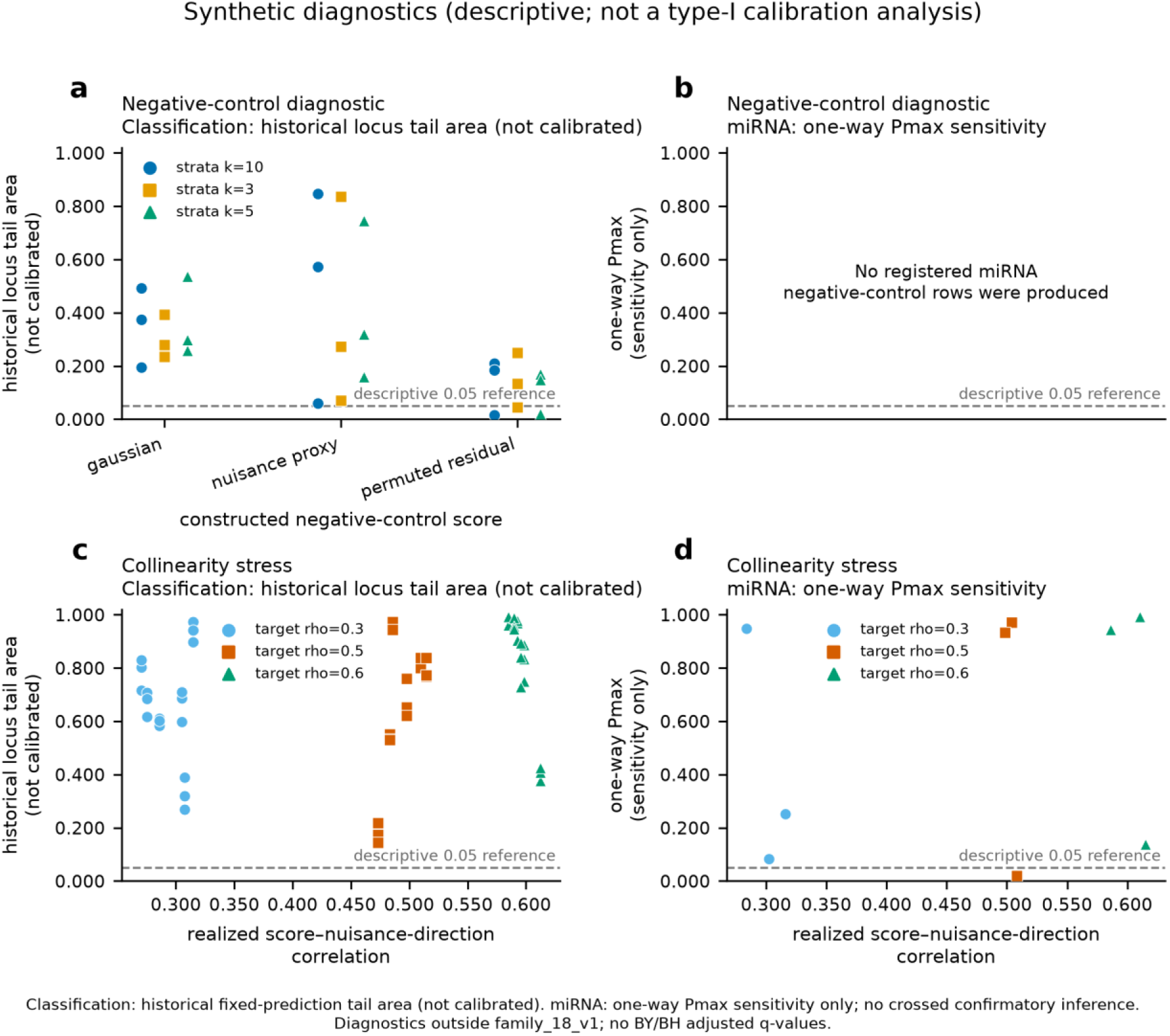
Diagnostic checks of negative-control and collinearity-stress implementations. Label-independent negative-control scores across declared nuisance-stratum settings are separated into (a) classification rows displaying historical fixed-prediction locus tail areas and (b) the registered miRNA panel; panel b records that no current miRNA negative-control rows were produced. Collinearity stress is separated into (c) classification rows and (d) miRNA rows. Across the collinearity panels, the maximum observed score-nuisance correlation was +0.6151, and 1 of 63 configured rows had a positive native delta and a role-appropriate displayed diagnostic tail area below 0.05. These diagnostics characterize the specified implementation settings; general calibration requires a separate evaluation (Supplementary Tables S3 and S14).

### Simple sequence baseline sensitivity

Fixed sequence composition provided an interpretable baseline comparison. The initial post-audit analysis added fixed mono-, di– and trinucleotide composition S to each measured baseline N under matched strict-inner fitting (Table 3; Table S15). Baseline gains were +0.0562 AUROC for CRISPRi minimal, +0.0205 with accessibility, and +0.0214/+0.0539 Spearman for HeLa/HEK293FT. Cas13 AUROC fell from 0.5897 to 0.5481. This initial restricted-grid result is recorded under its R7 identity, with later fits reported separately.

**Table 3.** Initial R7 measured and sequence composition baselines (restricted alpha grid)

| Member | n | Metric | Measured<br>baseline N | N plus<br>composition S | Sequence<br>increment |
| --- | --- | --- | --- | --- | --- |
| CRISPRi<br>minimal | 2961 | AUROC | 0.6426 | 0.6988 | +0.0562 |
| CRISPRi<br>accessibility | 2961 | AUROC | 0.7282 | 0.7487 | +0.0205 |
| Cas13 | 871 | AUROC | 0.5897 | 0.5481 | -0.0416 |
| HeLa | 2944 | Spearman | 0.4201 | 0.4415 | +0.0214 |
| HEK293FT | 2726 | Spearman | 0.3421 | 0.3960 | +0.0539 |

In the initial comparison, RNA-FM increments over N versus N+S were +0.0455/-0.0014 for CRISPRi minimal, +0.0189/-0.0007 with accessibility, –0.0105/-0.0022 for Cas13, +0.0001/– 0.0010 for HeLa and +0.0213/-0.0013 for HEK293FT. AlphaGenome, Enformer and Borzoi retained CRISPRi minimal increments over N+S of +0.0538, +0.0380 and +0.0486 (Fig. 4b,c; Table S15). All composition-baseline folds selected the grid maximum, alpha 100. A common-alpha diagnostic retained the main attenuation direction (Table S16), motivating a wider search over regularization. The later expanded-grid sensitivity retained all 18 cells and extended candidate alpha to 1,000,000. N+S selected 100 or 1000, never that upper bound (Table S17; Fig. S2). CRISPRi and HeLa estimates were unchanged. Cas13 N+S AUROC was 0.5532, still below N by 0.0365; HEK293FT N+S Spearman was 0.3927. RNA-FM increments over N+S became +0.0001 and +0.000044 for Cas13 and HEK293FT, respectively. The wider grid resolved boundary selection and identified which small increments were sensitive to regularization. Together with the retained positive CRISPRi increments from the DNA feature blocks, these results illustrate the value of testing both baseline content and fitting choices. The contrasts describe prediction under the specified probe and baseline; they do not quantify a causal proportion explained or establish absent information.

Together, these analyses show where sequence-model features add predictive value and which baseline or fitting choices alter that estimate. Matched members, explicit representations and recorded adjustment sets make those comparisons interpretable and reproducible within the evaluated assays.

### Practical guidance

The protocol turns these findings into a practical sequence of decisions: define the assay target and baseline, isolate downstream fitting, and report native-scale increments with their uncertainty and provenance (Box 1; Fig. 1a; Supplementary Methods; Box 2; Fig. 1b).

#### Box 1 Native scale model added last benchmarking protocol

1. Define the assay target. Record the read-out, evaluated unit and membership.
2. Match the biological baseline. Declare covariates and their sources; different adjustment sets define different conditional questions.
3. Fit and evaluate probes. Restrict adaptive fitting to outer-training rows and declare the inner preprocessing boundary.
4. Keep native scales. Report delta AUROC or delta cross-validated Spearman without pooling them.
5. State the dependence and inference status. Report one-way miRNA clustering sensitivities separately from joint crossed-cluster confirmation.
6. Describe resampling. State that fixed-prediction ranges condition on the fitted model; record per-arm valid fractions and reject arms below the declared threshold.
7. Compare simple sequence baselines. Report how model increments change on the same members when sequence information is added.
8. Justify practical size. Specify a task-native smallest effect of interest before assessing equivalence or negligibility.
9. Disclose provenance. Separate reused model inputs from new fitting, give file identities and report whether pretraining overlap was verified.

Table 3 compares measured-baseline and composition-augmented baseline performance.

The protocol provides a reproducible basis for comparing added predictive value under explicit assay and baseline definitions.

#### Box 2 Decision aid for interpreting added predictive information

Explicit read-out, baseline and fixed member? Define these together to specify the conditional predictive target.

Fixed evaluation rows and folds across the comparison? Report any membership changes separately from added information.

Outer-test isolation of adaptive fitting and explicit inner preprocessing? Document these boundaries and assess pretraining overlap separately.

Calibrated dependence-aware inference available? If unavailable, report point estimates and descriptive or sensitivity ranges with their stated roles.

Confirmatory inference intended? Justify the marginal-test and multiplicity assumptions; use descriptive reporting until they are supported.

Justified task-native SESOI? Assess practical size against that threshold; otherwise report the native increment without an equivalence claim.

Verified pretraining overlap? Use the provenance assessment to define the scope of independent evaluation.

Independent assay or cohort corroboration supports transportability only to the settings tested; mechanism requires a separate causal design.

Here, the decision aid supports native-scale predictive estimates under stated adjustments and identifies the additional evidence needed for mechanistic, practical-size or pretraining-independence conclusions.

## Discussion

The benchmark identifies positive sequence-model contributions and shows how biological baseline content shapes their interpretation. Similar augmented performance can coexist with different increments, as illustrated by the minimal and accessibility-adjusted lncRNA analyses. Composition, mature-guide identity and regularization each affected the residual increment. RNA-FM retained positive increments over the identity-supported domain baseline; adding composition reduced these increments, while positive AlphaGenome contributions remained in selected backgrounds. Cas13 composition lowered baseline performance, and broader regularization changed small RNA-FM signs. This heterogeneity makes baseline checking useful for identifying complementary predictive features rather than assuming that every added covariate strengthens every task. These findings complement existing RNA benchmarks [6, 10–12] and genomic-model limitations [13–15]. Tang et al. already compared pretrained DNA representations with one-hot and dinucleotide baselines [22]; our assay-matched composition sensitivity brings that principle into added-last comparisons with biological covariates.

The contribution is an operational framework that connects established predictive-value and benchmarking principles [16, 18, 19] to the choices encountered in ncRNA assays. Reporting the assay, membership, baseline, absolute performance, native increment and evaluation split together makes a model comparison easier to interpret and reproduce. The workflow also records representation provenance and the specific change tested by each sensitivity. It provides practical guidance using existing estimators; confirmatory calibration and practical-size decisions require their own justified inferential design and task-native smallest effect of interest.

Additional analyses broadened the examination of model and implementation choices. RiNALMo giga yielded a positive HEK293FT increment of +0.0238 (n = 2726; non-joint envelope +0.0123 to +0.0342), extending the observed pattern to another named RNA model (Table S18). Collinearity stress reached correlation +0.6151; 1/63 configured rows combined a positive increment with diagnostic tail area below 0.05 (Table S14). These checks describe the tested models and settings, with model-class generalization and calibration requiring broader evidence. The TargetScan analyses add a separate practical distinction: coverage selection and within-subset adjustment answer different questions from a same-member comparison of family and mature-miRNA aggregates.

## Limitations

The findings apply to conditional prediction in the evaluated assays. Mechanism, causal mediation and model-class capacity require broader experimental designs. Several features of the present analysis guide that next step. Resampling conditions on fixed predictions and omits training uncertainty; exchangeability of the observed-label permutation blocks remains unverified, and BY/BH transformations do not resolve it. The miRNA one-way arms leave joint crossed covariance unestimated. A prospectively justified task-native SESOI is also needed for equivalence or negligibility assessment. Pretraining overlap remains unverified, and upstream models were not rebuilt. For CRISPRi, labels pool cell lines whereas accessibility comes from K562; some original source-selection intermediates remain incomplete. The Cas13 source rule was independently recovered, although the original historical producer was not reconstructed. Community-port and official implementations are compared in Table S4. Guide hold-out addresses the declared partitions, with homology and broader seed-family transfer still to be evaluated. TargetScan sums could be refined by experimental isoform/site alignment, and extension beyond covered rows needs additional data. Expanded regularization resolved the tested boundary selection, while baseline optimality and the signs of very small increments remain sensitive to fitting choices. Later sensitivities were designed after preceding results and are descriptive. The incomplete assay/model matrix and finite diagnostics define the current scope; they also identify where independent cohorts and calibrated inference would most usefully extend the framework.

## Conclusions

Sequence-model features provided positive predictive increments in selected lncRNA growth or fitness and miRNA repression settings, with the size and direction depending on the baseline and fitting choices. The baseline-aware framework makes these contributions explicit by pairing native-scale increments with absolute performance, matched evaluation members, fold-local fitting and recorded provenance. Biological adjustment, simple sequence composition and mature-guide identity offer complementary ways to test what a model adds. This provides a practical foundation for selecting informative features and designing subsequent validation in ncRNA assays, with mechanistic interpretation and practical-size inference requiring their own evidence.

## Methods

### Members and labels

Each member fixed labels, covariates, chromosome folds and row identifiers across model sources. CRISPRi [7] marked a strict lncRNA hit in any of seven screened cell lines, not a K562-only outcome. The frozen table contained 3146 genes/188 strict hits; the common footprint contained 2961/174. Neighbor-only records were removed from positives but retained among screened non-strict hits; zero does not mean no growth phenotype. Accessibility adjustment used the same labels. For Cas13 [23], the historical 2308-gene/336-positive primary source set was independently recovered from both corrected Figshare v1 and final 2026 Table S1 using single explicit ENSG matches to GENCODE v49 lncRNA genes and excluding direct-ID strand discordance. Both reconstructed 14-column tables matched the frozen R6 table byte-for-byte. Versionless ENSG matching to E051 yielded 946/167; AlphaGenome availability defined 871/149. The earlier ID-only 2309/336 reconstruction is retained as a source-selection sensitivity; its one additional strand-discordant gene is outside E051 and both analysis intersections. MiRNA outcomes used the shared three-source footprint: HeLa 2944 and HEK293FT 2726. Two invalid mature-RNA intervals were excluded from the earlier HEK footprint and one retained row corrected. Missing required inputs were excluded without zero filling. The two-source coverage sensitivity retained those 946 E051 genes; 1363 genes/169 positives in the 2309-source population lacked the two-model common inputs and were excluded, not relabelled negative.

### Nuisance covariates

The minimal lncRNA baseline contained GTEx maximum expression, expression breadth and gene genomic length. Maximum expression was the maximum across 54 tissue-median TPM values; breadth was the fraction of these tissues with TPM at least 0.1, both rounded to four decimals in the input table. The length feature was log10 of genomic span (end minus start plus one), not mature transcript length. The expanded baseline added peak-derived K562 promoter DNase and ATAC measurements. GTEx and K562 features were not contemporaneous assay-matched measurements for every pooled CRISPRi label. MiRNA continuous covariates were target baseline expression, untranslated-region length, local AU content, distance to the untranslated-region end and site count. Seed type and transfected guide were categorical and one-hot encoded on outer-training rows, with unseen levels handled explicitly. The TargetScan gene-by-seed-family proxy was added only in the separately reported covered-subset sensitivity.

### Sequence definitions and models

LncRNA inputs used canonical mature transcripts, divided into non-overlapping near-equal windows within the model limit with complete coverage. Pooling weighted embeddings by nucleotide count; positional crops were sensitivities. MiRNA targets were mature-spliced 3′UTRs containing canonical sites; paired representations concatenated target, guide, product and absolute difference. DNA features were recorded AlphaGenome caches and Enformer/Borzoi community-port predictions [1–3]. Returned historical evidence matched all 8631 currently used AlphaGenome summary records. Matching raw arrays were available for 75 current miRNA requests (24 HeLa; 51 HEK293FT), with no additional current request recovered. For 68, separate original observation axes for both scorers were not retained. Recomputed aggregates agreed with the caches. Native session records supply user-authorization and execution context, while raw-response coverage remains partial. No new API or foundation-model inference was performed. Official Borzoi and TensorFlow Hub Enformer outputs supplied provenance sensitivities (Table S4). RNA-FM used released weights [5]; RiNALMo giga used an official checksum-locked checkpoint outside the primary family (Table S18). Model pretraining overlap remained unverified.

### Downstream fitting and added predictive value

For each member and model, delta-M was M(baseline plus model feature block) minus M(baseline), where M was AUROC for binary labels or cross-validated Spearman for continuous labels. These native scales remained separate. Within each outer fold, continuous baseline variables were transformed and standardised on training rows and categorical levels were learned there. Inner grouped cross-validation selected regularisation using baseline-only loss; the selected value was shared by baseline and augmented fits. A supervised probe compressed each model block to a scalar. The augmented model was trained on inner out-of-fold probe predictions and received the original covariates and that scalar. The probe was then refitted on outer-training rows for outer-test prediction. RNA-FM embeddings used 16 outer-training PCA components. In the reproduced primary implementation, baseline preprocessing and PCA were fitted once on the outer-training split before inner fitting. This preserves outer-test isolation, while inner-fold preprocessing shares the outer-training transformation. The post-audit analyses below additionally use strict inner preprocessing. The model blocks differ in dimension, track content and biological context. The contrast therefore evaluates each specified block and fitting pipeline. The polynomial baseline is a declared sensitivity. Cross-fitting provides methodological context [24]; the present target is predictive performance, distinct from the treatment and structural parameters addressed by that literature.

### Post audit sequence baseline comparison

The additional analysis was specified after the R6 audit, informed by earlier results. Its plan was locked before generating the new predictions as a post-audit sensitivity protocol. It retained all five task memberships, outcomes and chromosome folds and included every one of the 18 existing model cells. The sequence block S contained all ordered ACGU k-mers for k = 1, 2 and 3, with 84 normalized frequencies and no supervised feature selection. For lncRNA, frequencies were calculated within the same verified RNA-FM windows and averaged with nucleotide-length weights. For miRNA, composition of the same target crop and mature guide was concatenated with their elementwise product and absolute difference, giving 336 features. Invalid k-mers were excluded from frequency denominators and their counts recorded. These features match the mature-RNA support. DNA-model blocks additionally draw on their longer genomic context.

Both new baseline routes fitted preprocessing within each inner training split. Grouped validation selected alpha using classification log loss or continuous mean squared error; the selected baseline value was shared with its augmented fit. The initial grid was 0.1, 1, 10 and 100; the later sensitivity added 1000, 10000, 100000 and 1000000 without changing other rules. Model probes retained alpha 1, with training-local standardization and RNA-FM PCA16. Shared probe predictions supported paired N and N+S contrasts. Each analysis evaluated all 46 configurations by pooled outer-fold AUROC or Spearman. Comparisons within the new protocol isolate the added composition block. Historical-to-new differences additionally reflect stricter inner preprocessing. Fold metrics summarize the fitted splits descriptively. For TargetScan, exact full mature-sequence and seed matching defined the 1227/645-member sets; the terminal-U alternative was separate. Each set refitted six N/NS × none/family/guide backgrounds and all three models with fixed alpha 100, original chromosome groups and training-local preprocessing. Source weighted context++ rows were summed by gene with either seed-family or supported mature-miRNA identity, retaining genuine zeros and excluding missing coverage. The outputs are pooled point estimates and descriptive comparisons; no new P values, confidence intervals or equivalence thresholds were computed.

### Comparator tail areas, descriptive uncertainty and effect size

The configured lncRNA comparator ID was locus_label_permutation_within_outerfold_nuisance_quantile_fixed_crossfitted_predictions. Complete locus labels were permuted within blocks defined once from observed-label out-of-fold nuisance predictions, while the trained cross-fitted predictions were held fixed. This historical comparator is reported descriptively [25], because exchangeability of the observed-label blocks has not been established. The point estimator was fit once, and both bootstrap and null replicates evaluated the realised fixed cross-fitted predictions. The procedure conditions on the fitted model rather than refitting under each permuted label vector, leaving training-algorithm uncertainty outside its scope. Null-stratum count affects the comparator but not the point-estimation function. The unchanged point estimates in Table S19 follow directly from that computational separation.

Whiskers are the 2.5th and 97.5th percentiles of locus resampling with fitted cross-fitted predictions held fixed. They describe variation conditional on the fitted predictions. Confidence-interval coverage, power, minimum detectable effect, equivalence, detectability and claims of a small or absent effect require additional inferential assumptions and were not assessed. Locus resampling requested 1000 replicates. MiRNA ranges enveloped separate transcript– and guide-cluster arms. Separate transcript and guide wild-residual tail areas and their larger value describe one-way sensitivities. Joint crossed covariance was not estimated, so confirmatory miRNA P/q remained unavailable. A crossed pigeonhole bootstrap was also sensitivity-only. Each arm requested 1000 or its configured count, recorded valid and failed replicates, and failed closed below the declared valid fraction of 0.95.

Historical BY and BH transformations were calculated over the same declared 18-row accounting roster and retained only in supplementary audit material [26,27]. These transformations document the historical calculations. Confirmatory false-discovery-rate control would require valid marginal tests, which the fixed-prediction tail areas do not establish. The six unavailable miRNA inference fields were exact value-one sentinels in the machine schema; reader-facing tables show not assessed. One-way miRNA tail-area maxima, where retained for diagnostics, are separately labelled sensitivity summaries.

The analysis reports native-scale increments without a locked SESOI. Equivalence, minimum detectable effects, power and detectability were outside the scope of the descriptive ranges. The retired margin analysis is documented in Table S20. Primary folds were leave-one-chromosome-out, with leave-one-guide-out as a miRNA sensitivity. Analyses used seed 11. Negative controls, spike-ins and collinearity stress examine selected failure modes; a general calibration study remains a separate task.

### Reproducibility

The R6 snapshot was independently extracted and rerun on 12 September 2026; all 30 scoped downstream stages completed. Independent checks reproduced all 18 main point estimates and matched 4038 mapped figure-data fields. Additional R7/R8 fits reuse verified model inputs and are separately identified in tables and change records. The original reproduced implementation remains available alongside these extensions. Saved out-of-fold predictions support numerical recalculation of each sensitivity. Together, these records make the downstream calculations independently checkable using the recorded model inputs; upstream prediction generation and pretraining provenance remain separate components.

## Key Points

- A baseline-aware framework quantifies sequence-model contributions to lncRNA growth or fitness and miRNA repression using matched assay members.
- Native delta AUROC and delta cross-validated Spearman are reported alongside absolute baseline and augmented performance, making each increment interpretable.
- Sequence-model features retained positive lncRNA increments after accessibility adjustment, showing added predictive value under complementary biological baselines.
- Sequence composition, mature-miRNA feature identity and regularization reveal which model contributions persist across baseline choices and which depend on the fitting specification.

## Supplementary data

Supplementary Information and Supplementary Data 1 accompany this preprint.

## Funding

This work was supported by the National Natural Science Foundation of China (81500116 to Hong Wang; 81601449 to Rui Guo), the Jilin Science and Technology Development Program (YDZJ202301ZYTS092 to Rui Guo), the Wu Jieping Medical Foundation Clinical Research Special Fund (320.6750.2024-11-32 to Hong Wang), and the Jilin Tianhua Health Public Welfare Foundation (J2025JKJ064 to Hong Wang). The funders had no role in study design, data collection and analysis, the decision to publish, or preparation of the manuscript.

## Supporting information

Supplementary_Data_1_Main_Tables

## Acknowledgements

The authors thank the developers of the AlphaGenome, Enformer, Borzoi and RNA-FM models and of the TargetScan resource for making their tools and data publicly available. Use of AI-assisted tools in preparing this manuscript is disclosed in the declaration below.

## Declaration of generative AI and AI-assisted technologies in the writing process

We used Anthropic Claude through Claude Code for language editing, code assistance and technical auditing; OpenAI Codex for language editing, document preparation, code execution, clean-room reproduction and methodological review; and Google Gemini for academic literature search and citation-content verification. We also used Fable 5/claude-fable-5, GitHub Copilot, DeepSeek R1 and, in some rounds, Z.ai GLM to cross-audit code debugging. For the present revision, OpenAI Codex assisted with source-based corrections, numerical synchronization, code execution and preparation of the paragraph and table change ledger. The supplementary use log records this assistance. The authors retain responsibility for the wording, numerical results, citations and interpretation.

## Author contributions

H.W. conceived, designed and supervised the study, contributed to the bioinformatic analysis, critically reviewed the manuscript, and acquired funding. R.G. performed the bioinformatic and statistical analyses, contributed to study supervision, drafted the manuscript, and acquired funding. R.L. and W.S. carried out the literature search and prepared the figures. L.Y. provided supervision and interpreted the biological significance of the findings. D.M. supervised the bioinformatic and statistical analyses and the interpretation of the data.

## Conflict of interest

The authors declare that they have no competing interests.

## Ethics approval and consent to participate

Not applicable. This study analysed previously published, de-identified datasets and did not involve new human or animal subjects.

## Consent for publication

Not applicable.

## Data availability

This study reuses published data and model-derived inputs. The CRISPRi screen is Liu et al. [7] (PMID 27980086). Cas13 analyses use the corrected Figshare data package 10.6084/m9.figshare.30370171.v1 frozen in the R6 contract; the associated study was subsequently published as Essential lncRNAs in the human transcriptome [23]. The final 2026 supplementary Table S1 was independently obtained from Europe PMC. All 2576 explicit-ENSG endpoint labels, maximum essentiality counts and source-row counts agreed with Figshare v1. Annotation and formatting differences account for distinct workbook files; the R6 analysis retains its recorded data version. Uncorrected labels from the retracted 2024 Cell report [28] were not used. MiRNA data are from McGeary et al. [8], GEO GSE140217 and GSE140218; TargetScan is from Agarwal et al. [9]. K562 accessibility tracks are ENCODE ENCFF599DEH and ENCFF926KTI. Sequence inputs use GRCh38 and GENCODE under the mature-transcript and spliced-UTR policies described above. Hash-bound AlphaGenome caches, community-port Enformer and Borzoi scores and RNA-FM/RiNALMo embeddings were reused for downstream reproduction. Access and redistribution conditions differ by source. The retained June 2025 Output Terms restrict training for variant-effect prediction similar to AlphaGenome; they do not state a blanket ban on every supervised probe. Their applicability differs for users who accepted API Additional Terms. Native session records document user authorization for the September local execution phase. Provider permission and redistribution rights are separate from local authorization.

Accepted API Additional Terms, acceptance records and account-to-batch linkage were unavailable for review, leaving the applicable permission requirements and compliance status unresolved. Access to retained files therefore does not imply blanket redistribution rights for model outputs or weights. The Zenodo archive (https://doi.org/10.5281/zenodo.23393797) contains source manifests, aggregate results, final figure data and executable analysis and source-recovery code. Frozen fitting inputs are listed as external dependencies. Full downstream analysis requires access to those exact files; public availability has not been established for every frozen input. The supported, independently executed source-recovery contract applies single-explicit-ENSG matching and direct-ID strand concordance to either source workbook, yielding 2308 genes and 336 positives; both 14-column outputs have the same SHA256 as the frozen R6 primary table. The ID-only 2309-gene source is retained separately as a sensitivity. Both definitions preserve the 871/149 and 946/167 analysis cohorts. This independently recovers the source-selection rule and output. Reconstruction of the original historical producer and verification of pretraining overlap remain outside the completed recovery scope.

## Code availability

Code, input identities, source-specific recovery information and scoped execution records are available in the Zenodo release R10.20261015-v2 (https://doi.org/10.5281/zenodo.23393797), with separate instructions for reproducing the final figures from aggregate results. The supported route verifies and reuses eligible hash-bound model inputs and recomputes member tables, the 18-cell benchmark and sensitivities. The recovered Cas13 source contract is executable. Upstream reconstruction additionally depends on access, computational resources and source-specific terms. Some historical producers and source-selection intermediates remain incompletely reconstructed. The author-review documents and change ledger are separate from the public code and data release.

## Notes

https://zenodo.org/records/23393797

### Competing Interest Statement

The authors have declared no competing interest.

