## Supplementary_Information for "Evaluating the added value of sequence models in noncoding RNA assays: a baseline-aware framework"

This supplement documents the assay definitions, baseline comparisons and reproducibility checks underlying the framework. The CRISPRi outcome pools strict lncRNA hits across seven cell lines; K562 identifies the added accessibility covariates. The minimal length covariate is gene genomic span, and HEK task identifiers denote HEK293FT. Tables report freshly reproduced downstream calculations using recorded model inputs. Inference is labelled by its descriptive or sensitivity role.

### Supplementary Methods

TargetScan coverage denotes gene-by-seed-family records. The aggregate combines multiple mature miRNAs in 355 HeLa and 226 HEK293FT covered rows and serves as a family-level proxy. Guide-specific aggregate identity is examined in later sensitivities; experimental transcript/UTR-site alignment remains unavailable.

The primary implementation fits preprocessing and PCA on the outer-training split before inner fitting, preserving outer-test isolation. Separate post-audit analyses add strict inner preprocessing for the diagnostic and sequence-baseline comparisons.

The R8 analyses reported here were designed after examining the R6 and R7 results and locked before their own new fits. They report the full set of point estimates as descriptive sensitivities, including negative values; P values, confidence intervals, equivalence and model-class inference are outside their scope. Tables carried over from the R6 and R7 rounds retain their original numerical values under the identifiers used in this renumbered supplement; Fig. 4b,c also retains R7 values. The descriptions below distinguish the original results from subsequent sensitivities. N is the measured baseline, S fixed mature-RNA composition, and M the scalar model probe.

### Role-aware native-scale analysis tables

The tables pair task-native effects with the interpretation appropriate to each calculation. Classification tail areas are historical fixed-prediction comparator outputs, and whiskers are descriptive locus-resampling ranges conditional on the fitted predictions. They exclude training-algorithm uncertainty and are not calibrated confidence intervals. For crossed miRNA tasks, ranges envelop separate one-way procedures and Pmax summarizes sensitivity, without estimating joint crossed covariance. Unavailable inference is labelled not assessed. Equivalence, MDE, power, detectability and negligibility require additional design assumptions and are not assessed from these ranges. Native performance estimates, increments and descriptive range bounds are rounded once from stored values to four decimals throughout the text, tables and numerical figure annotations. Nonzero absolute values below 0.00005 use six decimals to retain their sign; full-precision estimates remain in the accompanying machine-readable tables. Axis ticks, configuration values and historical tail-area displays retain their separately stated scales.

**Table S1. Declared 18-cell benchmark accounting family. BY and BH columns retain historical arithmetic transformations without confirmatory error-rate control. For the six crossed-miRNA rows, exact 1/1 sentinels in the machine accounting record denote unavailable inference; the reader-facing display reports no inferential P or q.**

| Member | Model | n | Native metric/effect | Native delta | Role-aware inference display |
|---|---|---|---|---|---|
| e051_lncrna_cas13 | AG | 871 | auroc / delta_auroc | +0.0130 | descriptive fixed-OOF locus 95% range -0.0248 to +0.0475; historical fixed-prediction tail area=0.035; BY-18 adjusted value=0.244 (historical arithmetic only); BH-18 adjusted value=0.070 (assumption-sensitive display) |
| e051_lncrna_cas13 | Borzoi | 871 | auroc / delta_auroc | -0.0111 | descriptive fixed-OOF locus 95% range -0.0258 to +0.0035; historical fixed-prediction tail area=0.784; BY-18 adjusted value=1.000 (historical arithmetic only); BH-18 adjusted value=1.000 (assumption-sensitive display) |

| Member | Model | n | Native metric/effect | Native delta | Role-aware inference display |
|---|---|---|---|---|---|
| e051_lncrna_cas13 | Enformer | 871 | auroc / delta_auroc | -0.0161 | descriptive fixed-OOF locus 95% range -0.0293 to -0.0042; historical fixed-prediction tail area=0.896; BY-18 adjusted value=1.000 (historical arithmetic only); BH-18 adjusted value=1.000 (assumption-sensitive display) |
| e051_lncrna_cas13 | RNA-FM | 871 | auroc / delta_auroc | -0.0120 | descriptive fixed-OOF locus 95% range -0.0235 to -0.0006; historical fixed-prediction tail area=0.915; BY-18 adjusted value=1.000 (historical arithmetic only); BH-18 adjusted value=1.000 (assumption-sensitive display) |
| e051_lncrna_func | AG | 2961 | auroc / delta_auroc | +0.1125 | descriptive fixed-OOF locus 95% range +0.0680 to +0.1537; historical fixed-prediction tail area=<0.001; BY-18 adjusted value=0.009 (historical arithmetic only); BH-18 adjusted value=0.003 (assumption-sensitive display) |
| e051_lncrna_func | Borzoi | 2961 | auroc / delta_auroc | +0.0953 | descriptive fixed-OOF locus 95% range +0.0629 to +0.1293; historical fixed-prediction tail area=<0.001; BY-18 adjusted value=0.009 (historical arithmetic only); BH-18 adjusted value=0.003 (assumption-sensitive display) |
| e051_lncrna_func | Enformer | 2961 | auroc / delta_auroc | +0.0830 | descriptive fixed-OOF locus 95% range +0.0503 to +0.1153; historical fixed-prediction tail area=<0.001; BY-18 adjusted value=0.009 (historical arithmetic only); BH-18 adjusted value=0.003 (assumption-sensitive display) |
| e051_lncrna_func | RNA-FM | 2961 | auroc / delta_auroc | +0.0452 | descriptive fixed-OOF locus 95% range +0.0137 to +0.0780; historical fixed-prediction tail area=<0.001; BY-18 adjusted value=0.009 (historical arithmetic only); BH-18 adjusted value=0.003 (assumption-sensitive display) |
| e051_lncrna_func_acc | AG | 2961 | auroc / delta_auroc | +0.0252 | descriptive fixed-OOF locus 95% range +0.0094 to +0.0412; historical fixed-prediction tail area=<0.001; BY-18 adjusted value=0.009 (historical arithmetic only); BH-18 adjusted value=0.003 (assumption-sensitive display) |
| e051_lncrna_func_acc | Borzoi | 2961 | auroc / delta_auroc | +0.0246 | descriptive fixed-OOF locus 95% range +0.0101 to +0.0382; historical fixed-prediction tail area=<0.001; BY-18 adjusted value=0.009 (historical arithmetic only); BH-18 adjusted value=0.003 (assumption-sensitive display) |
| e051_lncrna_func_acc | Enformer | 2961 | auroc / delta_auroc | +0.0171 | descriptive fixed-OOF locus 95% range +0.0055 to +0.0282; historical fixed-prediction tail area=<0.001; BY-18 adjusted value=0.009 (historical arithmetic only); BH-18 adjusted value=0.003 (assumption-sensitive display) |
| e051_lncrna_func_acc | RNA-FM | 2961 | auroc / delta_auroc | +0.0184 | descriptive fixed-OOF locus 95% range +0.0040 to +0.0328; historical fixed-prediction tail area=0.006; BY-18 adjusted value=0.047 (historical arithmetic only); BH-18 adjusted value=0.013 (assumption-sensitive display) |

| Member | Model | n | Native metric/effect | Native delta | Role-aware inference display |
|---|---|---|---|---|---|
| e071_mirna_disrupt | AG | 2944 | cv_spearman / delta_cv_spearman | +0.0008 | non-joint one-way 95% resampling envelope -0.0005 to +0.0020; one-way Pmax sensitivity=0.033; crossed confirmatory P/q not assessed |
| e071_mirna_expr | AG | 2944 | cv_spearman / delta_cv_spearman | -0.0004 | non-joint one-way 95% resampling envelope -0.0013 to +0.0004; one-way Pmax sensitivity=0.735; crossed confirmatory P/q not assessed |
| e071_mirna_hek | RNA-FM | 2726 | cv_spearman / delta_cv_spearman | +0.0214 | non-joint one-way 95% resampling envelope +0.0107 to +0.0306; one-way Pmax sensitivity=<0.001; crossed confirmatory P/q not assessed |
| e071_mirna_hek_disrupt | AG | 2726 | cv_spearman / delta_cv_spearman | +0.0034 | non-joint one-way 95% resampling envelope -0.0013 to +0.0076; one-way Pmax sensitivity=0.040; crossed confirmatory P/q not assessed |
| e071_mirna_hek_expr | AG | 2726 | cv_spearman / delta_cv_spearman | -0.0008 | non-joint one-way 95% resampling envelope -0.0022 to +0.0010; one-way Pmax sensitivity=0.819; crossed confirmatory P/q not assessed |
| e071_mirna_hela | RNA-FM | 2944 | cv_spearman / delta_cv_spearman | +0.0002 | non-joint one-way 95% resampling envelope -0.0012 to +0.0015; one-way Pmax sensitivity=0.303; crossed confirmatory P/q not assessed |

**Table S2. Absolute baseline and model-augmented performance for the 18-cell family.** For every member-by-model cell, M(nuisance) is the cross-validated performance of that member's declared measured baseline. M(nuisance + model) adds a nested outer-training probe of the model feature block. Their difference, delta-M(native), remains on the native metric (AUROC for classification, cross-validated Spearman for continuous labels). This shows whether each increment sits on a strong or weak baseline. Rendered from the locked leaderboard.

| # | Member | Model | Native metric | M(nuisance) | M(nuisance + model) | delta-M (native) |
|---|---|---|---|---|---|---|
| 1 | e051_lncrna_cas13 | AG | AUROC | 0.5898 | 0.6028 | 0.0130 |
| 2 | e051_lncrna_cas13 | Borzoi | AUROC | 0.5898 | 0.5787 | -0.0111 |
| 3 | e051_lncrna_cas13 | Enformer | AUROC | 0.5898 | 0.5738 | -0.0161 |
| 4 | e051_lncrna_cas13 | RNA-FM | AUROC | 0.5898 | 0.5778 | -0.0120 |
| 5 | e051_lncrna_func | AG | AUROC | 0.6426 | 0.7551 | 0.1125 |
| 6 | e051_lncrna_func | Borzoi | AUROC | 0.6426 | 0.7378 | 0.0953 |
| 7 | e051_lncrna_func | Enformer | AUROC | 0.6426 | 0.7255 | 0.0830 |

| # | Member | Model | Native metric | M(nuisance) | M(nuisance + model) | delta-M (native) |
|---|---|---|---|---|---|---|
| 8 | e051_lncrna_func | RNA-FM | AUROC | 0.6426 | 0.6877 | 0.0452 |
| 9 | e051_lncrna_func_acc | AG | AUROC | 0.7282 | 0.7534 | 0.0252 |
| 10 | e051_lncrna_func_acc | Borzoi | AUROC | 0.7282 | 0.7528 | 0.0246 |
| 11 | e051_lncrna_func_acc | Enformer | AUROC | 0.7282 | 0.7454 | 0.0171 |
| 12 | e051_lncrna_func_acc | RNA-FM | AUROC | 0.7282 | 0.7467 | 0.0184 |
| 13 | e071_mirna_disrupt | AG | CV-Spearman | 0.4201 | 0.4208 | 0.0008 |
| 14 | e071_mirna_expr | AG | CV-Spearman | 0.4201 | 0.4197 | -0.0004 |
| 15 | e071_mirna_hek | RNA-FM | CV-Spearman | 0.3421 | 0.3635 | 0.0214 |
| 16 | e071_mirna_hek_disrupt | AG | CV-Spearman | 0.3421 | 0.3454 | 0.0034 |
| 17 | e071_mirna_hek_expr | AG | CV-Spearman | 0.3421 | 0.3413 | -0.0008 |
| 18 | e071_mirna_hela | RNA-FM | CV-Spearman | 0.4201 | 0.4203 | 0.0002 |

**Table S3. Synthetic protocol diagnostics and one-way dependence sensitivities.**

These analyses examine behavior on constructed inputs outside family_18_v1, without primary BY or BH-sensitivity adjusted q-values. Their role is to check specified implementations; establishing null exchangeability, type-I-error control, calibration or biological validity requires separate evaluation. ΔAUROC and ΔSpearman remain distinct native scales. Every displayed range and P-value summary below evaluates fixed cross-fitted out-of-fold predictions. Ranges condition on the fitted pipeline and exclude training-algorithm uncertainty. They are descriptive rather than confidence intervals. Requested/valid/failed counts are read directly from each analysis row.

### Table S3a. Synthetic positive-control diagnostic.

*ΔAUROC analyses*

| Analysis | ΔAUROC | Descriptive fixed-prediction locus 95% range | Historical fixed-prediction locus tail area (not calibrated) | Bootstrap n per arm; total requested/valid/failed | Null n per arm; total requested/valid/failed |
|---|---|---|---|---|---|
| e051_lncrna_func_acc; amplitude=0.2 | +0.0842 | +0.0610 to +0.1119 | <0.001 | 1000 per arm; 1000/1000/0 | 1000 per arm; 1000/1000/0 |
| e051_lncrna_func_acc; amplitude=0.6 | +0.2199 | +0.1876 to +0.2567 | <0.001 | 1000 per arm; 1000/1000/0 | 1000 per arm; 1000/1000/0 |

### Table S3b. Synthetic negative-control diagnostic.

*ΔAUROC analyses*

| Analysis | ΔAUROC | Descriptive fixed-prediction locus 95% range | Historical fixed-prediction locus tail area (not calibrated) | Bootstrap n per arm; total requested/valid/failed | Null n per arm; total requested/valid/failed |
|---|---|---|---|---|---|
| e051_lncrna_func_acc; gaussian; rep=0; strata=3 | +0.0012 | -0.0044 to +0.0086 | 0.393 | 200 per arm; 200/200/0 | 200 per arm; 200/200/0 |
| e051_lncrna_func_acc; gaussian; rep=0; strata=5 | +0.0012 | -0.0058 to +0.0085 | 0.537 | 200 per arm; 200/200/0 | 200 per arm; 200/200/0 |
| e051_lncrna_func_acc; gaussian; rep=0; strata=10 | +0.0012 | -0.0056 to +0.0083 | 0.493 | 200 per arm; 200/200/0 | 200 per arm; 200/200/0 |
| e051_lncrna_func_acc; gaussian; rep=1; strata=3 | +0.0023 | -0.0021 to +0.0065 | 0.279 | 200 per arm; 200/200/0 | 200 per arm; 200/200/0 |
| e051_lncrna_func_acc; gaussian; rep=1; strata=5 | +0.0023 | -0.0014 to +0.0078 | 0.259 | 200 per arm; 200/200/0 | 200 per arm; 200/200/0 |
| e051_lncrna_func_acc; gaussian; rep=1; strata=10 | +0.0023 | -0.0022 to +0.0062 | 0.373 | 200 per arm; 200/200/0 | 200 per arm; 200/200/0 |
| e051_lncrna_func_acc; gaussian; rep=2; strata=3 | +0.0020 | -0.0020 to +0.0066 | 0.234 | 200 per arm; 200/200/0 | 200 per arm; 200/200/0 |
| e051_lncrna_func_acc; gaussian; rep=2; strata=5 | +0.0020 | -0.0021 to +0.0068 | 0.299 | 200 per arm; 200/200/0 | 200 per arm; 200/200/0 |
| e051_lncrna_func_acc; gaussian; rep=2; strata=10 | +0.0020 | -0.0020 to +0.0064 | 0.194 | 200 per arm; 200/200/0 | 200 per arm; 200/200/0 |
| e051_lncrna_func_acc; nuisance_proxy; rep=0; strata=3 | +0.0050 | -0.0005 to +0.0101 | 0.070 | 200 per arm; 200/200/0 | 200 per arm; 200/200/0 |
| e051_lncrna_func_acc; nuisance_proxy; rep=0; strata=5 | +0.0050 | -0.0004 to +0.0105 | 0.159 | 200 per arm; 200/200/0 | 200 per arm; 200/200/0 |
| e051_lncrna_func_acc; nuisance_proxy; rep=0; strata=10 | +0.0050 | -0.0006 to +0.0107 | 0.060 | 200 per arm; 200/200/0 | 200 per arm; 200/200/0 |

| Analysis | ΔAUROC | Descriptive fixed-prediction locus 95% range | Historical fixed-prediction locus tail area (not calibrated) | Bootstrap n per arm; total requested/valid/failed | Null n per arm; total requested/valid/failed |
|---|---|---|---|---|---|
| e051_lncrna_func_acc; nuisance_proxy; rep=1; strata=3 | -0.0009 | -0.0031 to +0.0014 | 0.836 | 200 per arm; 200/200/0 | 200 per arm; 200/200/0 |
| e051_lncrna_func_acc; nuisance_proxy; rep=1; strata=5 | -0.0009 | -0.0035 to +0.0016 | 0.746 | 200 per arm; 200/200/0 | 200 per arm; 200/200/0 |
| e051_lncrna_func_acc; nuisance_proxy; rep=1; strata=10 | -0.0009 | -0.0040 to +0.0012 | 0.846 | 200 per arm; 200/200/0 | 200 per arm; 200/200/0 |
| e051_lncrna_func_acc; nuisance_proxy; rep=2; strata=3 | +0.0018 | -0.0058 to +0.0091 | 0.274 | 200 per arm; 200/200/0 | 200 per arm; 200/200/0 |
| e051_lncrna_func_acc; nuisance_proxy; rep=2; strata=5 | +0.0018 | -0.0052 to +0.0088 | 0.318 | 200 per arm; 200/200/0 | 200 per arm; 200/200/0 |
| e051_lncrna_func_acc; nuisance_proxy; rep=2; strata=10 | +0.0018 | -0.0057 to +0.0084 | 0.572 | 200 per arm; 200/200/0 | 200 per arm; 200/200/0 |
| e051_lncrna_func_acc; permuted_residual; rep=0; strata=3 | +0.0036 | -0.0003 to +0.0075 | 0.134 | 200 per arm; 200/200/0 | 200 per arm; 200/200/0 |
| e051_lncrna_func_acc; permuted_residual; rep=0; strata=5 | +0.0036 | +0.0002 to +0.0073 | 0.169 | 200 per arm; 200/200/0 | 200 per arm; 200/200/0 |
| e051_lncrna_func_acc; permuted_residual; rep=0; strata=10 | +0.0036 | +0.0003 to +0.0070 | 0.209 | 200 per arm; 200/200/0 | 200 per arm; 200/200/0 |
| e051_lncrna_func_acc; permuted_residual; rep=1; strata=3 | +0.0027 | -0.0022 to +0.0082 | 0.249 | 200 per arm; 200/200/0 | 200 per arm; 200/200/0 |
| e051_lncrna_func_acc; permuted_residual; rep=1; strata=5 | +0.0027 | -0.0019 to +0.0079 | 0.149 | 200 per arm; 200/200/0 | 200 per arm; 200/200/0 |
| e051_lncrna_func_acc; permuted_residual; rep=1; strata=10 | +0.0027 | -0.0021 to +0.0083 | 0.184 | 200 per arm; 200/200/0 | 200 per arm; 200/200/0 |
| e051_lncrna_func_acc; permuted_residual; rep=2; strata=3 | +0.0061 | +0.0008 to +0.0138 | 0.045 | 200 per arm; 200/200/0 | 200 per arm; 200/200/0 |
| e051_lncrna_func_acc; permuted_residual; rep=2; strata=5 | +0.0061 | +0.0001 to +0.0132 | 0.020 | 200 per arm; 200/200/0 | 200 per arm; 200/200/0 |
| e051_lncrna_func_acc; permuted_residual; rep=2; strata=10 | +0.0061 | +0.0007 to +0.0132 | 0.015 | 200 per arm; 200/200/0 | 200 per arm; 200/200/0 |

**Table S3c. miRNA transcript/guide one-way sensitivity decomposition.** These are extracted one-way arms evaluated on fixed cross-fitted out-of-fold predictions. The envelope and P-value maximum are sensitivity-only summaries, not joint crossed-cluster inference.

| Analysis | Native effect | Non-joint 95% resampling envelope (sensitivity only) | Transcript one-way 95% range; boot valid/failed | Guide one-way 95% range; boot valid/failed | Transcript wild P (sensitivity) | Guide wild P (sensitivity) | One-way Pmax (sensitivity only) |
|---|---|---|---|---|---|---|---|
| e071_mirna_hek · RNA-FM (ΔSpearman) | +0.0214 | +0.0107 to +0.0306 | +0.0107 to +0.0306; 1000/1000/0 | +0.0152 to +0.0280; 1000/1000/0 | <0.001 | <0.001 | <0.001 |
| e071_mirna_hela · RNA-FM (ΔSpearman) | +0.0002 | -0.0012 to +0.0015 | -0.0010 to +0.0013; 1000/1000/0 | -0.0012 to +0.0015; 1000/1000/0 | 0.274 | 0.303 | 0.303 |

**Table S3d. Collinear negative-control stress diagnostic.**

*ΔAUROC analyses*

| Analysis | ΔAUROC | Descriptive fixed-prediction locus 95% range | Historical fixed-prediction locus tail area (not calibrated) | Bootstrap n per arm; total requested/valid/failed | Null n per arm; total requested/valid/failed |
|---|---|---|---|---|---|
| e051_lncrna_func_acc; rho=0.3; rep=0; strata=3 | +0.0012 | -0.0046 to +0.0054 | 0.268 | 200 per arm; 200/200/0 | 1000 per arm; 1000/1000/0 |
| e051_lncrna_func_acc; rho=0.3; rep=0; strata=5 | +0.0012 | -0.0034 to +0.0064 | 0.389 | 200 per arm; 200/200/0 | 1000 per arm; 1000/1000/0 |
| e051_lncrna_func_acc; rho=0.3; rep=0; strata=10 | +0.0012 | -0.0039 to +0.0068 | 0.319 | 200 per arm; 200/200/0 | 1000 per arm; 1000/1000/0 |
| e051_lncrna_func_acc; rho=0.3; rep=1; strata=3 | -0.0004 | -0.0069 to +0.0080 | 0.597 | 200 per arm; 200/200/0 | 1000 per arm; 1000/1000/0 |
| e051_lncrna_func_acc; rho=0.3; rep=1; strata=5 | -0.0004 | -0.0084 to +0.0070 | 0.686 | 200 per arm; 200/200/0 | 1000 per arm; 1000/1000/0 |
| e051_lncrna_func_acc; rho=0.3; rep=1; strata=10 | -0.0004 | -0.0073 to +0.0065 | 0.710 | 200 per arm; 200/200/0 | 1000 per arm; 1000/1000/0 |
| e051_lncrna_func_acc; rho=0.3; rep=2; strata=3 | -0.0007 | -0.0029 to +0.0011 | 0.715 | 200 per arm; 200/200/0 | 1000 per arm; 1000/1000/0 |
| e051_lncrna_func_acc; rho=0.3; rep=2; strata=5 | -0.0007 | -0.0030 to +0.0015 | 0.803 | 200 per arm; 200/200/0 | 1000 per arm; 1000/1000/0 |
| e051_lncrna_func_acc; rho=0.3; rep=2; strata=10 | -0.0007 | -0.0029 to +0.0018 | 0.829 | 200 per arm; 200/200/0 | 1000 per arm; 1000/1000/0 |
| e051_lncrna_func_acc; rho=0.5; rep=0; strata=3 | +0.0019 | -0.0030 to +0.0072 | 0.174 | 200 per arm; 200/200/0 | 1000 per arm; 1000/1000/0 |
| e051_lncrna_func_acc; rho=0.5; rep=0; strata=5 | +0.0019 | -0.0032 to +0.0078 | 0.145 | 200 per arm; 200/200/0 | 1000 per arm; 1000/1000/0 |

| Analysis | ΔAUROC | Descriptive fixed-prediction locus 95% range | Historical fixed-prediction locus tail area (not calibrated) | Bootstrap n per arm; total requested/valid/failed | Null n per arm; total requested/valid/failed |
|---|---|---|---|---|---|
| e051_lncrna_func_acc; rho=0.5; rep=0; strata=10 | +0.0019 | -0.0032 to +0.0069 | 0.219 | 200 per arm; 200/200/0 | 1000 per arm; 1000/1000/0 |
| e051_lncrna_func_acc; rho=0.5; rep=1; strata=3 | -0.0018 | -0.0063 to +0.0029 | 0.823 | 200 per arm; 200/200/0 | 1000 per arm; 1000/1000/0 |
| e051_lncrna_func_acc; rho=0.5; rep=1; strata=5 | -0.0018 | -0.0066 to +0.0027 | 0.801 | 200 per arm; 200/200/0 | 1000 per arm; 1000/1000/0 |
| e051_lncrna_func_acc; rho=0.5; rep=1; strata=10 | -0.0018 | -0.0070 to +0.0024 | 0.839 | 200 per arm; 200/200/0 | 1000 per arm; 1000/1000/0 |
| e051_lncrna_func_acc; rho=0.5; rep=2; strata=3 | -0.0003 | -0.0069 to +0.0065 | 0.652 | 200 per arm; 200/200/0 | 1000 per arm; 1000/1000/0 |
| e051_lncrna_func_acc; rho=0.5; rep=2; strata=5 | -0.0003 | -0.0075 to +0.0071 | 0.620 | 200 per arm; 200/200/0 | 1000 per arm; 1000/1000/0 |
| e051_lncrna_func_acc; rho=0.5; rep=2; strata=10 | -0.0003 | -0.0072 to +0.0064 | 0.759 | 200 per arm; 200/200/0 | 1000 per arm; 1000/1000/0 |
| e051_lncrna_func_acc; rho=0.6; rep=0; strata=3 | +0.0013 | -0.0027 to +0.0051 | 0.377 | 200 per arm; 200/200/0 | 1000 per arm; 1000/1000/0 |
| e051_lncrna_func_acc; rho=0.6; rep=0; strata=5 | +0.0013 | -0.0026 to +0.0052 | 0.408 | 200 per arm; 200/200/0 | 1000 per arm; 1000/1000/0 |
| e051_lncrna_func_acc; rho=0.6; rep=0; strata=10 | +0.0013 | -0.0029 to +0.0058 | 0.425 | 200 per arm; 200/200/0 | 1000 per arm; 1000/1000/0 |
| e051_lncrna_func_acc; rho=0.6; rep=1; strata=3 | -0.0017 | -0.0038 to -0.000026 | 0.961 | 200 per arm; 200/200/0 | 1000 per arm; 1000/1000/0 |
| e051_lncrna_func_acc; rho=0.6; rep=1; strata=5 | -0.0017 | -0.0038 to +0.0004 | 0.992 | 200 per arm; 200/200/0 | 1000 per arm; 1000/1000/0 |
| e051_lncrna_func_acc; rho=0.6; rep=1; strata=10 | -0.0017 | -0.0037 to -0.0001 | 0.992 | 200 per arm; 200/200/0 | 1000 per arm; 1000/1000/0 |
| e051_lncrna_func_acc; rho=0.6; rep=2; strata=3 | -0.0017 | -0.0086 to +0.0037 | 0.750 | 200 per arm; 200/200/0 | 1000 per arm; 1000/1000/0 |
| e051_lncrna_func_acc; rho=0.6; rep=2; strata=5 | -0.0017 | -0.0075 to +0.0047 | 0.833 | 200 per arm; 200/200/0 | 1000 per arm; 1000/1000/0 |
| e051_lncrna_func_acc; rho=0.6; rep=2; strata=10 | -0.0017 | -0.0083 to +0.0043 | 0.887 | 200 per arm; 200/200/0 | 1000 per arm; 1000/1000/0 |
| e051_lncrna_func; rho=0.3; rep=0; strata=3 | -0.0067 | -0.0119 to -0.0008 | 0.973 | 200 per arm; 200/200/0 | 1000 per arm; 1000/1000/0 |
| e051_lncrna_func; rho=0.3; rep=0; strata=5 | -0.0067 | -0.0118 to -0.0018 | 0.941 | 200 per arm; 200/200/0 | 1000 per arm; 1000/1000/0 |
| e051_lncrna_func; rho=0.3; rep=0; strata=10 | -0.0067 | -0.0122 to -0.0018 | 0.898 | 200 per arm; 200/200/0 | 1000 per arm; 1000/1000/0 |
| e051_lncrna_func; rho=0.3; rep=1; strata=3 | -0.0113 | -0.0230 to +0.0011 | 0.708 | 200 per arm; 200/200/0 | 1000 per arm; 1000/1000/0 |
| e051_lncrna_func; rho=0.3; rep=1; strata=5 | -0.0113 | -0.0258 to +0.0018 | 0.683 | 200 per arm; 200/200/0 | 1000 per arm; 1000/1000/0 |
| e051_lncrna_func; rho=0.3; rep=1; strata=10 | -0.0113 | -0.0242 to +0.0004 | 0.617 | 200 per arm; 200/200/0 | 1000 per arm; 1000/1000/0 |

| Analysis | ΔAUROC | Descriptive fixed-prediction locus 95% range | Historical fixed-prediction locus tail area (not calibrated) | Bootstrap n per arm; total requested/valid/failed | Null n per arm; total requested/valid/failed |
|---|---|---|---|---|---|
| e051_lncrna_func; rho=0.3; rep=2; strata=3 | -0.0046 | -0.0114 to +0.0027 | 0.583 | 200 per arm; 200/200/0 | 1000 per arm; 1000/1000/0 |
| e051_lncrna_func; rho=0.3; rep=2; strata=5 | -0.0046 | -0.0121 to +0.0029 | 0.611 | 200 per arm; 200/200/0 | 1000 per arm; 1000/1000/0 |
| e051_lncrna_func; rho=0.3; rep=2; strata=10 | -0.0046 | -0.0117 to +0.0034 | 0.601 | 200 per arm; 200/200/0 | 1000 per arm; 1000/1000/0 |
| e051_lncrna_func; rho=0.5; rep=0; strata=3 | -0.0034 | -0.0074 to -0.0002 | 0.972 | 200 per arm; 200/200/0 | 1000 per arm; 1000/1000/0 |
| e051_lncrna_func; rho=0.5; rep=0; strata=5 | -0.0034 | -0.0068 to +0.0001 | 0.973 | 200 per arm; 200/200/0 | 1000 per arm; 1000/1000/0 |
| e051_lncrna_func; rho=0.5; rep=0; strata=10 | -0.0034 | -0.0073 to +0.0001 | 0.943 | 200 per arm; 200/200/0 | 1000 per arm; 1000/1000/0 |
| e051_lncrna_func; rho=0.5; rep=1; strata=3 | -0.0151 | -0.0299 to +0.0006 | 0.768 | 200 per arm; 200/200/0 | 1000 per arm; 1000/1000/0 |
| e051_lncrna_func; rho=0.5; rep=1; strata=5 | -0.0151 | -0.0301 to -0.0017 | 0.772 | 200 per arm; 200/200/0 | 1000 per arm; 1000/1000/0 |
| e051_lncrna_func; rho=0.5; rep=1; strata=10 | -0.0151 | -0.0342 to -0.0010 | 0.837 | 200 per arm; 200/200/0 | 1000 per arm; 1000/1000/0 |
| e051_lncrna_func; rho=0.5; rep=2; strata=3 | -0.0133 | -0.0280 to +0.0031 | 0.550 | 200 per arm; 200/200/0 | 1000 per arm; 1000/1000/0 |
| e051_lncrna_func; rho=0.5; rep=2; strata=5 | -0.0133 | -0.0297 to +0.0035 | 0.532 | 200 per arm; 200/200/0 | 1000 per arm; 1000/1000/0 |
| e051_lncrna_func; rho=0.5; rep=2; strata=10 | -0.0133 | -0.0290 to +0.0009 | 0.529 | 200 per arm; 200/200/0 | 1000 per arm; 1000/1000/0 |
| e051_lncrna_func; rho=0.6; rep=0; strata=3 | -0.0019 | -0.0041 to +0.0006 | 0.903 | 200 per arm; 200/200/0 | 1000 per arm; 1000/1000/0 |
| e051_lncrna_func; rho=0.6; rep=0; strata=5 | -0.0019 | -0.0041 to +0.0004 | 0.969 | 200 per arm; 200/200/0 | 1000 per arm; 1000/1000/0 |
| e051_lncrna_func; rho=0.6; rep=0; strata=10 | -0.0019 | -0.0042 to +0.0008 | 0.978 | 200 per arm; 200/200/0 | 1000 per arm; 1000/1000/0 |
| e051_lncrna_func; rho=0.6; rep=1; strata=3 | -0.0037 | -0.0069 to -0.0007 | 0.988 | 200 per arm; 200/200/0 | 1000 per arm; 1000/1000/0 |
| e051_lncrna_func; rho=0.6; rep=1; strata=5 | -0.0037 | -0.0075 to -0.0004 | 0.957 | 200 per arm; 200/200/0 | 1000 per arm; 1000/1000/0 |
| e051_lncrna_func; rho=0.6; rep=1; strata=10 | -0.0037 | -0.0069 to -0.0008 | 0.945 | 200 per arm; 200/200/0 | 1000 per arm; 1000/1000/0 |
| e051_lncrna_func; rho=0.6; rep=2; strata=3 | -0.0109 | -0.0213 to -0.0021 | 0.892 | 200 per arm; 200/200/0 | 1000 per arm; 1000/1000/0 |
| e051_lncrna_func; rho=0.6; rep=2; strata=5 | -0.0109 | -0.0202 to -0.0014 | 0.840 | 200 per arm; 200/200/0 | 1000 per arm; 1000/1000/0 |
| e051_lncrna_func; rho=0.6; rep=2; strata=10 | -0.0109 | -0.0227 to -0.0010 | 0.729 | 200 per arm; 200/200/0 | 1000 per arm; 1000/1000/0 |

*ΔSpearman analyses*

| Analysis | ΔSpearman | Non-joint one-way 95% resampling envelope (sensitivity only) | One-way transcript/guide Pmax (sensitivity only) | Bootstrap n per arm; total requested/valid/failed | Null n per arm; total requested/valid/failed |
|---|---|---|---|---|---|
| e071_mirna_hek_disrupt; rho=0.3; rep=0; strata=NOT_APPLICABLE | +0.0012 | -0.0010 to +0.0034 | 0.083 | 200 per arm; 400/400/0 | 1000 per arm; 2000/2000/0 |
| e071_mirna_hek_disrupt; rho=0.3; rep=1; strata=NOT_APPLICABLE | -0.0022 | -0.0046 to -0.0003 | 0.948 | 200 per arm; 400/400/0 | 1000 per arm; 2000/2000/0 |
| e071_mirna_hek_disrupt; rho=0.3; rep=2; strata=NOT_APPLICABLE | -0.0001 | -0.0028 to +0.0026 | 0.252 | 200 per arm; 400/400/0 | 1000 per arm; 2000/2000/0 |
| e071_mirna_hek_disrupt; rho=0.5; rep=0; strata=NOT_APPLICABLE | -0.0011 | -0.0024 to +0.0001 | 0.971 | 200 per arm; 400/400/0 | 1000 per arm; 2000/2000/0 |
| e071_mirna_hek_disrupt; rho=0.5; rep=1; strata=NOT_APPLICABLE | +0.0032 | +0.0003 to +0.0060 | 0.020 | 200 per arm; 400/400/0 | 1000 per arm; 2000/2000/0 |
| e071_mirna_hek_disrupt; rho=0.5; rep=2; strata=NOT_APPLICABLE | -0.0023 | -0.0046 to -0.0003 | 0.932 | 200 per arm; 400/400/0 | 1000 per arm; 2000/2000/0 |
| e071_mirna_hek_disrupt; rho=0.6; rep=0; strata=NOT_APPLICABLE | -0.0018 | -0.0034 to -0.0003 | 0.993 | 200 per arm; 400/400/0 | 1000 per arm; 2000/2000/0 |
| e071_mirna_hek_disrupt; rho=0.6; rep=1; strata=NOT_APPLICABLE | +0.0004 | -0.0030 to +0.0037 | 0.139 | 200 per arm; 400/400/0 | 1000 per arm; 2000/2000/0 |
| e071_mirna_hek_disrupt; rho=0.6; rep=2; strata=NOT_APPLICABLE | -0.0033 | -0.0064 to +0.0002 | 0.944 | 200 per arm; 400/400/0 | 1000 per arm; 2000/2000/0 |

For classification, the displayed historical tail area uses label permutation within outer-fold × nuisance-score blocks on fixed predictions. The locus-bootstrap whisker describes variation conditional on the fixed predictions. Both displays have descriptive rather than calibrated confirmatory roles. For miRNA, the display combines separate transcript and guide wild-residual sensitivities, their P-value maximum and a non-joint one-way envelope. Crossed confirmatory inference remains unavailable. The numerical p_confirmatory=1 unavailable-inference sentinel is never displayed as a P value. Equivalence, TOST, MDE, power, detectability and negligibility are not assessed from these ranges.

**Supplementary Note 2. Official model-port concordance and Enformer sensitivity.**

The locked benchmark uses cached community-port Enformer and Borzoi scores. To check port fidelity, official inference was run on all 3146 E051 lncRNA loci. Borzoi used the official Baskerville hound_predbed.py script and Enformer the DeepMind TensorFlow Hub model. The outputs were compared with the cached scores feature by feature. The feature-level concordance metrics are reported without a post hoc qualitative threshold in Tables S4a-b. The official Enformer features were then carried through the same downstream estimator, with resampling conditional on fixed cross-fitted predictions, as a sensitivity (Table S4c). The locked primary leaderboard and its historical BY-18 arithmetic are retained alongside these checks. Replacement-family BY-18 and BH-18 calculations describe counterfactual substitutions, with the latter additionally dependent on its assumptions.

**Table S4.** Official model-port concordance (Enformer, Borzoi) and the official-Enformer downstream sensitivity.

**Table S4a. Enformer official (DeepMind TensorFlow Hub) versus cached community-port features, n = 3146 loci.**

| Feature | Pearson r | Spearman rho | Top-decile overlap | RMSE |
|---|---|---|---|---|
| ENF_DNASE_mean | 0.9823 | 0.9907 | 0.9175 | 0.5872 |
| ENF_DNASE_max | 0.9886 | 0.9860 | 0.9492 | 5.198 |
| ENF_ATAC_mean | 0.9676 | 0.9736 | 0.8857 | 3.828 |
| ENF_ATAC_max | 0.9764 | 0.9769 | 0.8603 | 11.48 |
| ENF_CAGE_mean | 0.9938 | 0.9802 | 0.9460 | 1.394 |

| Feature | Pearson r | Spearman rho | Top-decile overlap | RMSE |
|---|---|---|---|---|
| ENF_CAGE_max | 0.9859 | 0.9756 | 0.9111 | 38.68 |
| ENF_H3K27ac_mean | 0.9630 | 0.9705 | 0.8476 | 6.146 |
| ENF_H3K27ac_max | 0.9335 | 0.9432 | 0.6349 | 24.6 |

**Table S4b. Borzoi official (Baskerville hound_predbed.py) versus cached community-port features, n = 3146 loci.**

| Feature | Pearson r | Spearman rho | Top-decile overlap | RMSE |
|---|---|---|---|---|
| BZ_RNA_SEQ_mean | 1.0000 | 1.0000 | 1.0000 | 0.0001093 |
| BZ_RNA_SEQ_max | 1.0000 | 1.0000 | 1.0000 | 0.003854 |
| BZ_DNASE_mean | 1.0000 | 1.0000 | 1.0000 | 0.0002105 |
| BZ_DNASE_max | 1.0000 | 1.0000 | 1.0000 | 0.005331 |
| BZ_ATAC_mean | 1.0000 | 1.0000 | 1.0000 | 0.0001372 |
| BZ_ATAC_max | 1.0000 | 1.0000 | 1.0000 | 0.004335 |
| BZ_CAGE_mean | 1.0000 | 1.0000 | 1.0000 | 0.0001565 |
| BZ_CAGE_max | 1.0000 | 1.0000 | 0.9968 | 0.0175 |
| BZ_H3K27ac_mean | 1.0000 | 1.0000 | 1.0000 | 0.001664 |
| BZ_H3K27ac_max | 1.0000 | 1.0000 | 1.0000 | 0.013 |

**Table S4c. Official-Enformer downstream sensitivity through the same downstream estimator.** Only the three lncRNA Enformer cells are recomputed; the displayed named q-values are the counterfactual historical BY-18 and BH-18 arithmetic values after substituting these cells into the unchanged 18-row roster.

| Cell | Native scale | Locked-port Δ | Official Δ | Official 2.5th–97.5th percentile descriptive conditional range | Historical-comparator one-sided locus-permutation P (not calibrated) | Replacement-family BY-18 q (primary method) | Replacement-family BH-18 q (sensitivity) |
|---|---|---|---|---|---|---|---|
| lncRNA-Cas13 | delta_auroc | -0.0161 | -0.0037 | -0.0319 to +0.0257 | 0.2178 | 1.0000 | 0.3920 |
| lncRNA-func | delta_auroc | +0.0830 | +0.0907 | +0.0559 to +0.1262 | 0.000999 | 0.0090 | 0.0026 |
| lncRNA-func-acc | delta_auroc | +0.0171 | +0.0181 | +0.0068 to +0.0301 | 0.000999 | 0.0090 | 0.0026 |

Feature-level concordance and downstream native-scale sensitivity answer different questions and are therefore reported separately. The official-Enformer cells provide an additional sensitivity outside the locked primary family. Replacement-family calculations use counterfactual substitution within the unchanged 18-row roster, while the primary result retains its original identity.

Whiskers are the 2.5th and 97.5th percentiles of locus resampling with fitted cross-fitted predictions held fixed. They describe conditional variation. Confidence-interval coverage, power, minimum detectable effect, equivalence, detectability and small-or-absent-effect assessment require additional inferential support and were not evaluated.

Complete locus labels were permuted within blocks defined once from observed-label out-of-fold nuisance predictions, while the trained cross-fitted predictions were held fixed. The resulting historical comparator summarizes that construction descriptively; its marginal P values are not calibrated confirmatory tests.

Primary multiplicity adjustment used Benjamini-Yekutieli over the locked 18-row family. This procedure accommodates arbitrary dependence across hypotheses only when the marginal P values are valid or super-uniform; that marginal-validity condition is not established by the adjustment.

Benjamini-Hochberg adjustment over the same locked 18-row family was retained only as an assumption-sensitive secondary analysis because no independence or positive-regression-dependence argument was established.

Point estimates, descriptive ranges, locus-permutation P values and named multiplicity outputs are reported with their distinct roles. They characterize conditional prediction; cis/trans mechanism is a separate causal question.

**Table S5. Cas13 source-population and two-source coverage sensitivity.**

Additional provenance records enabled recovery of the direct-ID strand-concordance rule. An independent source entry accepts a single explicit ENSG token uniquely mapped to a GENCODE v49 lncRNA gene, excludes all direct-ID source/GTF strand discordances, and aggregates any-positive status and the maximum essential-cell-line count at gene level. Executed separately on corrected Figshare v1 and final 2026 Table S1, it recovered the historical 2308 genes/336 positives. Both complete 14-column TSVs matched the frozen R6 SHA256. The sole excluded discordant record is Hum_XLOC_002176 (S1A Excel row 217; ENSG00000260805.4), with source strand + and GTF strand −. The source/GTF annotation conflict remains unresolved. The previous ID-only reconstruction of 2309/336 is retained as a source-selection sensitivity. The discordant gene lies outside E051, preserving both the 871 and 946 analysis cohorts and their fitted inputs. Independent execution recovered the source rule and output; the original historical producer was not rerun. The retained two-source and covariate intersection is 946/167, all within E051. The four-model intersection remains 871/149. The added 75/18 are evaluated using the available Enformer and Borzoi inputs; AlphaGenome and RNA-FM coverage remains unavailable. Composition was computed from 116 mature-RNA windows for the added genes without RNA-model inference. Both Enformer and Borzoi use their retained community-port feature blocks; the separate TensorFlow Hub and Hound official outputs belong to the earlier provenance sensitivities and were not substituted here. All original 871 labels, N, S, folds and two retained DNA feature matrices were verified unchanged. Both new baselines and both augmented routes were refitted with strict inner processing and the eight-value grid. The six configurations describe coverage sensitivity within the same study.

**Table S5a. Cas13 ID-only source-selection sensitivity and coverage populations.**

This table retains the previously analysed ID-only 2309-gene source population and its original denominators. The subsequently recovered strand-concordant historical primary source contains 2308 genes/336 positives; its 946-gene complement is 1362, whereas the ID-only sensitivity complement remains 1363. The 871/149 and 946/167 fitted cohorts and all reported model results are identical under these two source definitions. These alternative eligibility definitions support a direct check of source-selection sensitivity within the study.

| Population | n | Positive | Negative | Positive fraction | Fraction of 2309 source |
|---|---|---|---|---|---|
| Source_2309 | 2309 | 336 | 1973 | 0.1455 | 1.0000 |
| All_models_871 | 871 | 149 | 722 | 0.1711 | 0.3772 |
| Two_DNA_946 | 946 | 167 | 779 | 0.1765 | 0.4097 |
| New_75 | 75 | 18 | 57 | 0.2400 | 0.0325 |
| Source_without_two_DNA_1363 | 1363 | 169 | 1194 | 0.1240 | 0.5903 |
| Source_without_all_models_1438 | 1438 | 187 | 1251 | 0.1300 | 0.6228 |

**Table S5b. 946-member baseline performance.**

| Task | n | Metric | N | N+S | S increment |
|---|---|---|---|---|---|
| cas13_dna_coverage | 946 | AUROC | 0.5784 | 0.5481 | -0.0303 |

**Table S5c. Both 946-member model comparisons.**

| Task | Model | n | Metric | N | N+M | N+S | N+S+M | M given N | M given N+S |
|---|---|---|---|---|---|---|---|---|---|
| cas13_dna_coverage | Enformer | 946 | AUROC | 0.5784 | 0.5675 | 0.5481 | 0.5445 | -0.0109 | -0.0036 |
| cas13_dna_coverage | Borzoi | 946 | AUROC | 0.5784 | 0.5765 | 0.5481 | 0.5479 | -0.0019 | -0.0002 |

**Table S5d. Original and extended coverage comparisons.**

| Population | Model | n | N | N+M | N+S | N+S+M | M given N | M given N+S |
|---|---|---|---|---|---|---|---|---|
| all_model871 | Enformer | 871 | 0.5897 | 0.5753 | 0.5532 | 0.5510 | -0.0145 | -0.0022 |
| two_DNA946 | Enformer | 946 | 0.5784 | 0.5675 | 0.5481 | 0.5445 | -0.0109 | -0.0036 |
| all_model871 | Borzoi | 871 | 0.5897 | 0.5818 | 0.5532 | 0.5533 | -0.0079 | +0.0001 |
| two_DNA946 | Borzoi | 946 | 0.5784 | 0.5765 | 0.5481 | 0.5479 | -0.0019 | -0.0002 |

All N outer folds selected alpha 10; N+S selected 1000 in 23 folds and 100 in one, below the upper grid limit. The two populations differ in both evaluation members and training distributions. The Borzoi increment over N+S changes from +0.0001 at 871 to -0.0002 at 946. This small sign change indicates sensitivity to coverage. The composition baseline performs worse in both populations. These overlapping-population comparisons describe fitting and coverage sensitivity, leaving absence and equivalence unassessed.

### Table S6. Historical TargetScan coverage and family-aggregate adjustment.

The three tracks separate the full-cohort reference, coverage-subset selection and within-subset family-aggregate adjustment. These historical fits are separate from the fixed-alpha exact-identity refits in Table 2 and Table S7.

#### Table S6a. RNA-FM increments in the three historical fitting tracks.

| Cohort | Track | Track n / full n | Baseline rho | Augmented rho | RNA-FM delta | Non-joint one-way 95% envelope | One-way Pmax |
|---|---|---|---|---|---|---|---|
| HEK293FT | A. Full cohort | 2726 / 2726 | 0.3421 | 0.3635 | +0.0214 | +0.0107 to +0.0306 | <0.001 |
| HEK293FT | B. Covered subset | 705 / 2726 | 0.3914 | 0.3972 | +0.0058 | +0.0002 to +0.0151 | 0.022 |
| HEK293FT | C. Covered subset + family aggregate | 705 / 2726 | 0.4412 | 0.4423 | +0.0012 | -0.0035 to +0.0099 | 0.186 |
| HeLa | A. Full cohort | 2944 / 2944 | 0.4201 | 0.4203 | +0.0002 | -0.0012 to +0.0015 | 0.303 |
| HeLa | B. Covered subset | 1227 / 2944 | 0.4250 | 0.4301 | +0.0052 | +0.0004 to +0.0102 | 0.002 |
| HeLa | C. Covered subset + family aggregate | 1227 / 2944 | 0.4592 | 0.4562 | -0.0029 | -0.0051 to -0.0007 | 0.999 |

A denotes the full cohort with the original nuisance covariates; B denotes the TargetScan-covered subset with the same nuisance covariates; C denotes the same covered subset after adding the family aggregate. Baseline rho is cross-validated Spearman correlation without RNA-FM; augmented rho adds RNA-FM. Each RNA-FM delta is augmented rho minus baseline rho.

Ranges are non-joint one-way 95% resampling envelopes for individual RNA-FM increments, and Pmax values summarize one-way sensitivity. Crossed confirmatory P/q is not assessed. Table S6b reports descriptive differences between increments, separately from these track-specific summaries.

The family feature sums weighted context++ source rows by gene and seed family, potentially combining mature miRNAs. It represents a gene-by-family proxy; experimental guide-specific total repression is a different quantity.

### Table S6b. Descriptive contrasts between the RNA-FM increments in Table S6a.

| Cohort | Covered n / full n | Covered-subset contrast B - A | Within-subset adjustment C - B |
|---|---|---|---|
| HEK293FT | 705 / 2726 | -0.0155 | -0.0047 |
| HeLa | 1227 / 2944 | +0.0050 | -0.0081 |

The covered-subset contrast is B - A; the within-subset adjustment contrast is C - B. The first captures changes in both population and fit, while the second compares increments within the same covered subset. Both are descriptive comparisons rather than causal effects.

Differences were calculated from unrounded estimates; subtraction of displayed values may differ in the final digit because each value is rounded independently. These are point contrasts without resampled inference. The track-specific Pmax values in Table S6a apply to individual increments, separately from the contrasts.

### Table S7. Exact-identity TargetScan analyses.

#### Table S7a. Exact-identity TargetScan baselines.

| Set | n | Background | Baseline rho | Domain gain versus none |
|---|---|---|---|---|
| hela_exact | 1227 | N__none | 0.3873 | +0.0000 |
| hela_exact | 1227 | N__family | 0.4140 | +0.0267 |
| hela_exact | 1227 | N__guide | 0.4799 | +0.0926 |
| hela_exact | 1227 | NS__none | 0.4199 | +0.0000 |
| hela_exact | 1227 | NS__family | 0.4417 | +0.0218 |
| hela_exact | 1227 | NS__guide | 0.4843 | +0.0643 |
| hek_exact | 645 | N__none | 0.3777 | +0.0000 |
| hek_exact | 645 | N__family | 0.4284 | +0.0507 |
| hek_exact | 645 | N__guide | 0.4358 | +0.0581 |
| hek_exact | 645 | NS__none | 0.3922 | +0.0000 |
| hek_exact | 645 | NS__family | 0.4386 | +0.0465 |
| hek_exact | 645 | NS__guide | 0.4463 | +0.0541 |

#### Table S7b. All 36 exact-identity model comparisons.

| Set | n | Background | Model | Baseline rho | Augmented rho | Model increment |
|---|---|---|---|---|---|---|
| hela_exact | 1227 | N__none | AlphaGenome_disrupt | 0.3873 | 0.3876 | +0.0003 |
| hela_exact | 1227 | N__none | AlphaGenome_expr | 0.3873 | 0.3862 | -0.0011 |
| hela_exact | 1227 | N__none | RNA-FM | 0.3873 | 0.4179 | +0.0306 |
| hela_exact | 1227 | N__family | AlphaGenome_disrupt | 0.4140 | 0.4142 | +0.0002 |
| hela_exact | 1227 | N__family | AlphaGenome_expr | 0.4140 | 0.4131 | -0.0009 |
| hela_exact | 1227 | N__family | RNA-FM | 0.4140 | 0.4439 | +0.0299 |
| hela_exact | 1227 | N__guide | AlphaGenome_disrupt | 0.4799 | 0.4799 | +0.000005 |
| hela_exact | 1227 | N__guide | AlphaGenome_expr | 0.4799 | 0.4795 | -0.0005 |

| Set | n | Background | Model | Baseline rho | Augmented rho | Model increment |
|---|---|---|---|---|---|---|
| hela_exact | 1227 | N__guide | RNA-FM | 0.4799 | 0.4969 | +0.0170 |
| hela_exact | 1227 | NS__none | AlphaGenome_disrupt | 0.4199 | 0.4194 | -0.0005 |
| hela_exact | 1227 | NS__none | AlphaGenome_expr | 0.4199 | 0.4207 | +0.0008 |
| hela_exact | 1227 | NS__none | RNA-FM | 0.4199 | 0.4206 | +0.0007 |
| hela_exact | 1227 | NS__family | AlphaGenome_disrupt | 0.4417 | 0.4414 | -0.0004 |
| hela_exact | 1227 | NS__family | AlphaGenome_expr | 0.4417 | 0.4426 | +0.0009 |
| hela_exact | 1227 | NS__family | RNA-FM | 0.4417 | 0.4420 | +0.0003 |
| hela_exact | 1227 | NS__guide | AlphaGenome_disrupt | 0.4843 | 0.4840 | -0.0003 |
| hela_exact | 1227 | NS__guide | AlphaGenome_expr | 0.4843 | 0.4858 | +0.0015 |
| hela_exact | 1227 | NS__guide | RNA-FM | 0.4843 | 0.4857 | +0.0014 |
| hek_exact | 645 | N__none | AlphaGenome_disrupt | 0.3777 | 0.3833 | +0.0056 |
| hek_exact | 645 | N__none | AlphaGenome_expr | 0.3777 | 0.3792 | +0.0015 |
| hek_exact | 645 | N__none | RNA-FM | 0.3777 | 0.3989 | +0.0212 |
| hek_exact | 645 | N__family | AlphaGenome_disrupt | 0.4284 | 0.4334 | +0.0051 |
| hek_exact | 645 | N__family | AlphaGenome_expr | 0.4284 | 0.4300 | +0.0017 |
| hek_exact | 645 | N__family | RNA-FM | 0.4284 | 0.4392 | +0.0108 |
| hek_exact | 645 | N__guide | AlphaGenome_disrupt | 0.4358 | 0.4404 | +0.0046 |
| hek_exact | 645 | N__guide | AlphaGenome_expr | 0.4358 | 0.4383 | +0.0025 |
| hek_exact | 645 | N__guide | RNA-FM | 0.4358 | 0.4434 | +0.0076 |
| hek_exact | 645 | NS__none | AlphaGenome_disrupt | 0.3922 | 0.3986 | +0.0064 |
| hek_exact | 645 | NS__none | AlphaGenome_expr | 0.3922 | 0.3936 | +0.0015 |
| hek_exact | 645 | NS__none | RNA-FM | 0.3922 | 0.3883 | -0.0038 |
| hek_exact | 645 | NS__family | AlphaGenome_disrupt | 0.4386 | 0.4441 | +0.0055 |
| hek_exact | 645 | NS__family | AlphaGenome_expr | 0.4386 | 0.4408 | +0.0022 |
| hek_exact | 645 | NS__family | RNA-FM | 0.4386 | 0.4362 | -0.0024 |
| hek_exact | 645 | NS__guide | AlphaGenome_disrupt | 0.4463 | 0.4514 | +0.0051 |
| hek_exact | 645 | NS__guide | AlphaGenome_expr | 0.4463 | 0.4487 | +0.0024 |
| hek_exact | 645 | NS__guide | RNA-FM | 0.4463 | 0.4425 | -0.0038 |

**Table S8. Terminal-U mapping assumption analyses.**

**Table S8a. Terminal-U mapping assumption baselines.**

| Set | n | Background | Baseline rho | Domain gain versus none |
|---|---|---|---|---|
| hek_tailU_proxy | 705 | N__none | 0.3804 | +0.0000 |
| hek_tailU_proxy | 705 | N__family | 0.4377 | +0.0573 |
| hek_tailU_proxy | 705 | N__guide | 0.4485 | +0.0682 |
| hek_tailU_proxy | 705 | NS__none | 0.3919 | +0.0000 |
| hek_tailU_proxy | 705 | NS__family | 0.4420 | +0.0501 |
| hek_tailU_proxy | 705 | NS__guide | 0.4478 | +0.0559 |

**Table S8b. All 18 terminal-U assumption model comparisons.**

| Set | n | Background | Model | Baseline rho | Augmented rho | Model increment |
|---|---|---|---|---|---|---|
| hek_tailU_proxy | 705 | N__none | AlphaGenome_disrupt | 0.3804 | 0.3858 | +0.0055 |
| hek_tailU_proxy | 705 | N__none | AlphaGenome_expr | 0.3804 | 0.3789 | -0.0014 |
| hek_tailU_proxy | 705 | N__none | RNA-FM | 0.3804 | 0.4054 | +0.0250 |
| hek_tailU_proxy | 705 | N__family | AlphaGenome_disrupt | 0.4377 | 0.4416 | +0.0040 |
| hek_tailU_proxy | 705 | N__family | AlphaGenome_expr | 0.4377 | 0.4357 | -0.0020 |
| hek_tailU_proxy | 705 | N__family | RNA-FM | 0.4377 | 0.4502 | +0.0125 |
| hek_tailU_proxy | 705 | N__guide | AlphaGenome_disrupt | 0.4485 | 0.4521 | +0.0035 |
| hek_tailU_proxy | 705 | N__guide | AlphaGenome_expr | 0.4485 | 0.4496 | +0.0011 |
| hek_tailU_proxy | 705 | N__guide | RNA-FM | 0.4485 | 0.4569 | +0.0084 |
| hek_tailU_proxy | 705 | NS__none | AlphaGenome_disrupt | 0.3919 | 0.3965 | +0.0046 |
| hek_tailU_proxy | 705 | NS__none | AlphaGenome_expr | 0.3919 | 0.3925 | +0.0006 |
| hek_tailU_proxy | 705 | NS__none | RNA-FM | 0.3919 | 0.3900 | -0.0019 |
| hek_tailU_proxy | 705 | NS__family | AlphaGenome_disrupt | 0.4420 | 0.4442 | +0.0022 |
| hek_tailU_proxy | 705 | NS__family | AlphaGenome_expr | 0.4420 | 0.4420 | -0.000009 |
| hek_tailU_proxy | 705 | NS__family | RNA-FM | 0.4420 | 0.4408 | -0.0012 |
| hek_tailU_proxy | 705 | NS__guide | AlphaGenome_disrupt | 0.4478 | 0.4501 | +0.0023 |
| hek_tailU_proxy | 705 | NS__guide | AlphaGenome_expr | 0.4478 | 0.4503 | +0.0025 |
| hek_tailU_proxy | 705 | NS__guide | RNA-FM | 0.4478 | 0.4459 | -0.0020 |

### TargetScan identity and fitting details

F sums human weighted context++ source rows by gene and seed family; G sums rows by gene and one identity-supported mature miRNA. Both may span source transcripts/sites and represent gene-level source aggregates. Experimental isoform-specific total repression and a performance upper bound are separate quantities. Exact identity requires full T-to-Unormalized mature sequence and nt2-8 seed agreement with a unique human entry. The resulting sets are HeLa 1227 rows/16 guides and HEK293FT 645 rows/11 guides. The HEK miR-133a experimental sequence has one additional terminal U relative to the compatible source entry. Retaining its 60 covered rows gives a separate 705-row/12-guide sensitivity to that mapping assumption. Non-human lsy-6 has no human counterpart. Genuine numeric zeros are retained; absent coverage is not replaced with zero.

Each set used the same members across six backgrounds (N or NS, each with no domain feature, F or G) and all three models. Baseline/meta alpha was fixed to 100, with inner-train transformations, 16 RNA-FM PCs and scalar-probe alpha 1. Twenty-three original chromosome outer groups and five grouped inner splits were used. Each covered subset was refitted to obtain its own OOF scores. All 72 baseline/augmented configurations, 18 baseline values and 54 model increments are retained. Pooled Spearman differences within each set use identical members. Comparing 645 with 705 additionally changes membership, so the terminal-U sensitivity addresses mapping and coverage together. Results are point estimates without P/CI calculations. Full protocol, source hashes, excluded IDs, 1242 fold descriptions and OOF predictions are retained under analysis/targetscan.

**Table S9. miRNA leave-one-guide-out sensitivity. One-way envelopes and crossed-pigeonhole ranges describe sensitivity under distinct dependence constructions. Their signs may differ, which informs interpretation of the held-out-guide estimates. Neither construction supplies calibrated joint confirmation, and crossed-cluster confirmatory P and multiplicity results remain unavailable.**

| Member | Model | Fold scheme | Native delta Spearman | Non-joint sensitivity display |
|---|---|---|---|---|
| e071_mirna_disrupt | AG | guide_loo(n=17) | +0.0010 | non-joint one-way 95% resampling envelope -0.0013 to +0.0031; one-way Pmax sensitivity=0.100; crossed confirmatory P/q not assessed; crossed-pigeonhole 95% sensitivity range -0.0021 to +0.0049; not a confirmatory interval |
| e071_mirna_expr | AG | guide_loo(n=17) | -0.0005 | non-joint one-way 95% resampling envelope -0.0018 to +0.0008; one-way Pmax sensitivity=0.645; crossed confirmatory P/q not assessed; crossed-pigeonhole 95% sensitivity range -0.0029 to +0.0018; not a confirmatory interval |
| e071_mirna_hek | RNA-FM | guide_loo(n=12) | +0.0153 | non-joint one-way 95% resampling envelope +0.0014 to +0.0378; one-way Pmax sensitivity=0.012; crossed confirmatory P/q not assessed; crossed-pigeonhole 95% sensitivity range -0.0002 to +0.0385; not a confirmatory interval |
| e071_mirna_hek_disrupt | AG | guide_loo(n=12) | +0.0045 | non-joint one-way 95% resampling envelope +0.0009 to +0.0085; one-way Pmax sensitivity=0.004; crossed confirmatory P/q not assessed; crossed-pigeonhole 95% sensitivity range -0.0022 to +0.0116; not a confirmatory interval |
| e071_mirna_hek_expr | AG | guide_loo(n=12) | -0.0020 | non-joint one-way 95% resampling envelope -0.0038 to -0.0002; one-way Pmax sensitivity=0.947; crossed confirmatory P/q not assessed; crossed-pigeonhole 95% sensitivity range -0.0048 to +0.0003; not a confirmatory interval |
| e071_mirna_hela | RNA-FM | guide_loo(n=17) | +0.0002 | non-joint one-way 95% resampling envelope -0.0142 to +0.0144; one-way Pmax sensitivity=0.419; crossed confirmatory P/q not assessed; crossed-pigeonhole 95% sensitivity range -0.0143 to +0.0145; not a confirmatory interval |

**Table S10. TargetScan seed-family assignment and guide/family LOO identity check.**

- hek: 12 guides, 12 mapped, 12 distinct families; guide-LOO and family-LOO are identical partitions.
- hela: 17 guides, 16 mapped, 17 distinct families; guide-LOO and family-LOO are identical partitions.

| Cohort | miRNA | miRBase ID | Seed family | Seed (2-8) | Mapped |
|---|---|---|---|---|---|
| hek | mir122 | hsa-miR-122-5p | miR-122-5p | GGAGUGU | 1 |
| hek | mir133 | hsa-miR-133a-3p.1 | miR-133a-3p.1 | UGGUCCC | 1 |
| hek | mir138 | hsa-miR-138-5p | miR-138-5p | GCUGGUG | 1 |
| hek | mir145 | hsa-miR-145-5p | miR-145-5p | UCCAGUU | 1 |
| hek | mir184 | hsa-miR-184 | miR-184 | GGACGGA | 1 |
| hek | mir190a | hsa-miR-190a-5p | miR-190-5p | GAUAUGU | 1 |
| hek | mir200b | hsa-miR-200b-3p | miR-200bc-3p/429 | AAUACUG | 1 |
| hek | mir216a | hsa-miR-216a-5p | miR-216a-5p | AAUCUCA | 1 |
| hek | mir217 | hsa-miR-217 | miR-217 | ACUGCAU | 1 |
| hek | mir219a | hsa-miR-219a-5p | miR-219-5p | GAUUGUC | 1 |
| hek | mir375 | hsa-miR-375 | miR-375 | UUGUUCG | 1 |
| hek | mir451a | hsa-miR-451a | miR-451 | AACCGUU | 1 |
| hela | let7 | hsa-let-7a-5p | let-7-5p/98-5p | GAGGUAG | 1 |
| hela | lsy6 | unmapped | OWN_FAMILY:lsy6 | NA | 0 |
| hela | mir1 | hsa-miR-1-3p | miR-1-3p/206 | GGAAUGU | 1 |
| hela | mir124 | hsa-miR-124-3p.1 | miR-124-3p.1 | AAGGCAC | 1 |
| hela | mir137 | hsa-miR-137 | miR-137 | UAUUGCU | 1 |
| hela | mir139 | hsa-miR-139-5p | miR-139-5p | CUACAGU | 1 |
| hela | mir143 | hsa-miR-143-3p | miR-143-3p | GAGAUGA | 1 |
| hela | mir144 | hsa-miR-144-3p | miR-144-3p | ACAGUAU | 1 |
| hela | mir153 | hsa-miR-153-3p | miR-153-3p | UGCAUAG | 1 |
| hela | mir155 | hsa-miR-155-5p | miR-155-5p | UAAUGCU | 1 |
| hela | mir182 | hsa-miR-182-5p | miR-182-5p | UUGGCAA | 1 |
| hela | mir199a | hsa-miR-199a-5p | miR-199-5p | CCAGUGU | 1 |
| hela | mir204 | hsa-miR-204-5p | miR-204-5p/211-5p | UCCCUUU | 1 |
| hela | mir205 | hsa-miR-205-5p | miR-205-5p | CCUUCAU | 1 |
| hela | mir216b | hsa-miR-216b-5p | miR-216b-5p | AAUCUCU | 1 |
| hela | mir223 | hsa-miR-223-3p | miR-223-3p | GUCAGUU | 1 |
| hela | mir7 | hsa-miR-7-5p | miR-7-5p | GGAAGAC | 1 |

**Table S11. Deterministic producer-path consistency check of the primary outer-training PCA point path.**

| Member | Native primary delta | Recomputed outer-PCA delta | Difference | Tolerance | Pass |
|---|---|---|---|---|---|
| e051_lncrna_cas13 | -0.0120 | -0.0120 | +0.0000 | 1.0e-10 | yes |
| e051_lncrna_func | +0.0452 | +0.0452 | +0.0000 | 1.0e-10 | yes |
| e051_lncrna_func_acc | +0.0184 | +0.0184 | +0.0000 | 1.0e-10 | yes |
| e071_mirna_hek | +0.0214 | +0.0214 | +0.0000 | 1.0e-10 | yes |
| e071_mirna_hela | +0.0002 | +0.0002 | +0.0000 | 1.0e-10 | yes |

**Table S12. Outer-test-isolated RNA-FM PCA-k point-estimate sensitivity. This exploratory sweep reports how the point estimate changes with representation dimension.**

| Member | k | n | Native metric/effect | Native delta | Role |
|---|---|---|---|---|---|
| e051_lncrna_cas13 | 3 | 871 | auroc / delta_auroc | -0.0129 | point estimate only |
| e051_lncrna_cas13 | 8 | 871 | auroc / delta_auroc | +0.0040 | point estimate only |
| e051_lncrna_cas13 | 10 | 871 | auroc / delta_auroc | -0.0038 | point estimate only |
| e051_lncrna_cas13 | 16 | 871 | auroc / delta_auroc | -0.0120 | point estimate only |
| e051_lncrna_func | 3 | 2961 | auroc / delta_auroc | +0.0188 | point estimate only |
| e051_lncrna_func | 8 | 2961 | auroc / delta_auroc | +0.0322 | point estimate only |
| e051_lncrna_func | 10 | 2961 | auroc / delta_auroc | +0.0393 | point estimate only |
| e051_lncrna_func | 16 | 2961 | auroc / delta_auroc | +0.0452 | point estimate only |
| e051_lncrna_func_acc | 3 | 2961 | auroc / delta_auroc | +0.0134 | point estimate only |
| e051_lncrna_func_acc | 8 | 2961 | auroc / delta_auroc | +0.0113 | point estimate only |
| e051_lncrna_func_acc | 10 | 2961 | auroc / delta_auroc | +0.0146 | point estimate only |
| e051_lncrna_func_acc | 16 | 2961 | auroc / delta_auroc | +0.0184 | point estimate only |
| e071_mirna_hek | 3 | 2726 | cv_spearman / delta_cv_spearman | +0.0123 | point estimate only |
| e071_mirna_hek | 8 | 2726 | cv_spearman / delta_cv_spearman | +0.0110 | point estimate only |
| e071_mirna_hek | 10 | 2726 | cv_spearman / delta_cv_spearman | +0.0104 | point estimate only |
| e071_mirna_hek | 16 | 2726 | cv_spearman / delta_cv_spearman | +0.0214 | point estimate only |
| e071_mirna_hela | 3 | 2944 | cv_spearman / delta_cv_spearman | -0.0009 | point estimate only |
| e071_mirna_hela | 8 | 2944 | cv_spearman / delta_cv_spearman | +0.0012 | point estimate only |
| e071_mirna_hela | 10 | 2944 | cv_spearman / delta_cv_spearman | +0.0026 | point estimate only |
| e071_mirna_hela | 16 | 2944 | cv_spearman / delta_cv_spearman | +0.0002 | point estimate only |

### Table S13. Primary RNA-FM path and downstream sensitivity analyses.

#### S13a. Primary RNA-FM analysis path. These rows reproduce the primary estimates for comparison with the sensitivities below.

| Member | Native delta | Role-aware inference display | Feature array |
|---|---|---|---|
| e051_lncrna_cas13 | -0.0120 | descriptive fixed-OOF locus 95% range -0.0235 to -0.0006; historical fixed-prediction tail area=0.915 | emb |
| e051_lncrna_func | +0.0452 | descriptive fixed-OOF locus 95% range +0.0137 to +0.0780; historical fixed-prediction tail area=<0.001 | emb |
| e051_lncrna_func_acc | +0.0184 | descriptive fixed-OOF locus 95% range +0.0040 to +0.0328; historical fixed-prediction tail area=0.006 | emb |
| e071_mirna_hek | +0.0214 | non-joint one-way 95% resampling envelope +0.0107 to +0.0306; one-way Pmax sensitivity=<0.001; crossed confirmatory P/q not assessed | paired_features |
| e071_mirna_hela | +0.0002 | non-joint one-way 95% resampling envelope -0.0012 to +0.0015; one-way Pmax sensitivity=0.303; crossed confirmatory P/q not assessed | paired_features |

#### S13b. Degree-2 nuisance sensitivity, reported separately from the primary family.

| Member/model | Linear-primary native delta | Degree-2 native delta | Shift | Degree-2 role-aware inference |
|---|---|---|---|---|
| e051_lncrna_func_acc / AG | +0.0252 | +0.0232 | -0.0020 | descriptive fixed-OOF locus 95% range +0.0083 to +0.0387; historical fixed-prediction tail area=<0.001 |
| e071_mirna_hek / RNA-FM | +0.0214 | +0.0171 | -0.0043 | non-joint one-way 95% resampling envelope +0.0080 to +0.0247; one-way Pmax sensitivity=<0.001; crossed confirmatory P/q not assessed |

#### S13c. Input-representation sensitivity, reported as point estimates.

| Member | Representation | n | Native effect | Native delta | Source array hash |
|---|---|---|---|---|---|
| e051_lncrna_cas13 | canonical_full_window_pool | 871 | delta_auroc | -0.0120 | 4cbed3fbeaf1… |
| e051_lncrna_cas13 | crop_3p_1022 | 871 | delta_auroc | -0.0089 | 2b0e63843d33… |
| e051_lncrna_cas13 | crop_5p_1022 | 871 | delta_auroc | +0.0060 | 2b0e63843d33… |
| e051_lncrna_cas13 | crop_center_1022 | 871 | delta_auroc | +0.0093 | 2b0e63843d33… |
| e051_lncrna_func | canonical_full_window_pool | 2961 | delta_auroc | +0.0452 | 4cbed3fbeaf1… |
| e051_lncrna_func | crop_3p_1022 | 2961 | delta_auroc | +0.0299 | 2b0e63843d33… |

| Member | Representation | n | Native effect | Native delta | Source array hash |
|---|---|---|---|---|---|
| e051_lncrna_func | crop_5p_1022 | 2961 | delta_auroc | +0.0443 | 2b0e63843d33… |
| e051_lncrna_func | crop_center_1022 | 2961 | delta_auroc | +0.0363 | 2b0e63843d33… |
| e051_lncrna_func_acc | canonical_full_window_pool | 2961 | delta_auroc | +0.0184 | 4cbed3fbeaf1… |
| e051_lncrna_func_acc | crop_3p_1022 | 2961 | delta_auroc | +0.0180 | 2b0e63843d33… |
| e051_lncrna_func_acc | crop_5p_1022 | 2961 | delta_auroc | +0.0138 | 2b0e63843d33… |
| e051_lncrna_func_acc | crop_center_1022 | 2961 | delta_auroc | +0.0184 | 2b0e63843d33… |
| e071_mirna_hek | mature_target_only_sensitivity | 2726 | delta_cv_spearman | +0.0306 | 13aeddff0833… |
| e071_mirna_hek | mature_target_plus_experimental_guide_primary | 2726 | delta_cv_spearman | +0.0214 | 13aeddff0833… |
| e071_mirna_hela | mature_target_only_sensitivity | 2944 | delta_cv_spearman | +0.0056 | 9f9d825ab2e1… |
| e071_mirna_hela | mature_target_plus_experimental_guide_primary | 2944 | delta_cv_spearman | +0.0002 | 9f9d825ab2e1… |

**Table S14. Collinear label-independent negative-control stress diagnostic. Classification rows report historical fixed-prediction locus tail areas; the crossed-miRNA row reports one-way Pmax sensitivity. These outputs describe the tested stress settings and are not calibrated tests or estimates of type-I error.**

| Member | Cells | Corr(score, nuisance direction) | Native delta range | Smallest role-appropriate display P among positive deltas |
|---|---|---|---|---|
| e051_lncrna_func | 27 | +0.2753 to +0.5954 | -0.0151 to -0.0019 | not applicable (no positive delta) |
| e051_lncrna_func_acc | 27 | +0.2706 to +0.6123 | -0.0018 to +0.0019 | 0.145 |
| e071_mirna_hek_disrupt | 9 | +0.2840 to +0.6151 | -0.0033 to +0.0032 | 0.020 |

Across the 63 registered diagnostic cells, 1 row had a positive native delta and role-appropriate display P below 0.05. The count summarizes the observed diagnostic behavior within these settings.

**Table S15. Initial R7 restricted-grid model increments after adding sequence composition.**

| Task and model | n | N | N plus M | N plus S | N plus S plus M | M given N | M given N plus S |
|---|---|---|---|---|---|---|---|
| e051_lncrna_func / AG | 2961 | 0.6426 | 0.7551 | 0.6988 | 0.7526 | +0.1125 | +0.0538 |
| e051_lncrna_func / Enformer | 2961 | 0.6426 | 0.7255 | 0.6988 | 0.7368 | +0.0830 | +0.0380 |
| e051_lncrna_func / Borzoi | 2961 | 0.6426 | 0.7377 | 0.6988 | 0.7475 | +0.0951 | +0.0486 |
| e051_lncrna_func / RNA-FM | 2961 | 0.6426 | 0.6880 | 0.6988 | 0.6974 | +0.0455 | -0.0014 |
| e051_lncrna_func_acc / AG | 2961 | 0.7282 | 0.7535 | 0.7487 | 0.7596 | +0.0252 | +0.0108 |
| e051_lncrna_func_acc / Enformer | 2961 | 0.7282 | 0.7454 | 0.7487 | 0.7552 | +0.0171 | +0.0065 |
| e051_lncrna_func_acc / Borzoi | 2961 | 0.7282 | 0.7528 | 0.7487 | 0.7616 | +0.0245 | +0.0129 |
| e051_lncrna_func_acc / RNA-FM | 2961 | 0.7282 | 0.7471 | 0.7487 | 0.7480 | +0.0189 | -0.0007 |
| e051_lncrna_cas13 / AG | 871 | 0.5897 | 0.6012 | 0.5481 | 0.5577 | +0.0115 | +0.0095 |
| e051_lncrna_cas13 / Enformer | 871 | 0.5897 | 0.5753 | 0.5481 | 0.5440 | -0.0145 | -0.0041 |
| e051_lncrna_cas13 / Borzoi | 871 | 0.5897 | 0.5818 | 0.5481 | 0.5453 | -0.0079 | -0.0029 |
| e051_lncrna_cas13 / RNA-FM | 871 | 0.5897 | 0.5792 | 0.5481 | 0.5460 | -0.0105 | -0.0022 |
| e071_mirna_hela / AlphaGenome_disrupt | 2944 | 0.4201 | 0.4208 | 0.4415 | 0.4420 | +0.0008 | +0.0005 |
| e071_mirna_hela / AlphaGenome_expr | 2944 | 0.4201 | 0.4197 | 0.4415 | 0.4407 | -0.0004 | -0.0008 |
| e071_mirna_hela / RNA-FM | 2944 | 0.4201 | 0.4202 | 0.4415 | 0.4405 | +0.0001 | -0.0010 |
| e071_mirna_hek / AlphaGenome_disrupt | 2726 | 0.3421 | 0.3454 | 0.3960 | 0.3997 | +0.0034 | +0.0037 |
| e071_mirna_hek / AlphaGenome_expr | 2726 | 0.3421 | 0.3413 | 0.3960 | 0.3959 | -0.0008 | -0.0001 |
| e071_mirna_hek / RNA-FM | 2726 | 0.3421 | 0.3634 | 0.3960 | 0.3947 | +0.0213 | -0.0013 |

All entries report point estimates from the matched new protocol, without P values or intervals.

**Table S16. R7 common alpha diagnostic.**

Separately locked after inspecting the primary extension; alpha is fixed to 100 in both routes.

| Task and model | n | N | N plus M | N plus S | N plus S plus M | M given N | M given N plus S |
|---|---|---|---|---|---|---|---|
| e051_lncrna_func / AG | 2961 | 0.6400 | 0.7589 | 0.6988 | 0.7526 | +0.1189 | +0.0538 |
| e051_lncrna_func / Enformer | 2961 | 0.6400 | 0.7292 | 0.6988 | 0.7368 | +0.0892 | +0.0380 |
| e051_lncrna_func / Borzoi | 2961 | 0.6400 | 0.7414 | 0.6988 | 0.7475 | +0.1014 | +0.0486 |

| Task and model | n | N | N plus M | N plus S | N plus S plus M | M given N | M given N plus S |
|---|---|---|---|---|---|---|---|
| e051_lncrna_func / RNA-FM | 2961 | 0.6400 | 0.6893 | 0.6988 | 0.6974 | +0.0493 | -0.0014 |
| e051_lncrna_func_acc / AG | 2961 | 0.7330 | 0.7579 | 0.7487 | 0.7596 | +0.0249 | +0.0108 |
| e051_lncrna_func_acc / Enformer | 2961 | 0.7330 | 0.7511 | 0.7487 | 0.7552 | +0.0180 | +0.0065 |
| e051_lncrna_func_acc / Borzoi | 2961 | 0.7330 | 0.7574 | 0.7487 | 0.7616 | +0.0244 | +0.0129 |
| e051_lncrna_func_acc / RNA-FM | 2961 | 0.7330 | 0.7497 | 0.7487 | 0.7480 | +0.0167 | -0.0007 |
| e051_lncrna_cas13 / AG | 871 | 0.5817 | 0.6020 | 0.5481 | 0.5577 | +0.0203 | +0.0095 |
| e051_lncrna_cas13 / Enformer | 871 | 0.5817 | 0.5648 | 0.5481 | 0.5440 | -0.0169 | -0.0041 |
| e051_lncrna_cas13 / Borzoi | 871 | 0.5817 | 0.5764 | 0.5481 | 0.5453 | -0.0053 | -0.0029 |
| e051_lncrna_cas13 / RNA-FM | 871 | 0.5817 | 0.5716 | 0.5481 | 0.5460 | -0.0101 | -0.0022 |
| e071_mirna_hela / AlphaGenome_disrupt | 2944 | 0.4090 | 0.4099 | 0.4415 | 0.4420 | +0.0008 | +0.0005 |
| e071_mirna_hela / AlphaGenome_expr | 2944 | 0.4090 | 0.4089 | 0.4415 | 0.4407 | -0.0001 | -0.0008 |
| e071_mirna_hela / RNA-FM | 2944 | 0.4090 | 0.4160 | 0.4415 | 0.4405 | +0.0070 | -0.0010 |
| e071_mirna_hek / AlphaGenome_disrupt | 2726 | 0.3344 | 0.3389 | 0.3960 | 0.3997 | +0.0045 | +0.0037 |
| e071_mirna_hek / AlphaGenome_expr | 2726 | 0.3344 | 0.3337 | 0.3960 | 0.3959 | -0.0006 | -0.0001 |
| e071_mirna_hek / RNA-FM | 2726 | 0.3344 | 0.3623 | 0.3960 | 0.3947 | +0.0279 | -0.0013 |

### Table S17. Expanded regularization-grid analyses.

### Table S17a. Expanded-grid baseline performance.

| Task | n | Metric | N | N+S | S increment |
|---|---|---|---|---|---|
| e051_lncrna_func | 2961 | AUROC | 0.6426 | 0.6988 | +0.0562 |
| e051_lncrna_func_acc | 2961 | AUROC | 0.7282 | 0.7487 | +0.0205 |
| e051_lncrna_cas13 | 871 | AUROC | 0.5897 | 0.5532 | -0.0365 |
| e071_mirna_hela | 2944 | Spearman | 0.4201 | 0.4415 | +0.0214 |
| e071_mirna_hek | 2726 | Spearman | 0.3421 | 0.3927 | +0.0506 |

### Table S17b. All 18 expanded-grid model comparisons.

| Task | Model | n | Metric | N | N+M | N+S | N+S+M | M given N | M given N+S |
|---|---|---|---|---|---|---|---|---|---|
| e051_lncrna_func | AG | 2961 | AUROC | 0.6426 | 0.7551 | 0.6988 | 0.7526 | +0.1125 | +0.0538 |
| e051_lncrna_func | Enformer | 2961 | AUROC | 0.6426 | 0.7255 | 0.6988 | 0.7368 | +0.0830 | +0.0380 |
| e051_lncrna_func | Borzoi | 2961 | AUROC | 0.6426 | 0.7377 | 0.6988 | 0.7475 | +0.0951 | +0.0486 |

| Task | Model | n | Metric | N | N+M | N+S | N+S+M | M given N | M given N+S |
|---|---|---|---|---|---|---|---|---|---|
| e051_lncrna_func | RNA-FM | 2961 | AUROC | 0.6426 | 0.6880 | 0.6988 | 0.6974 | +0.0455 | -0.0014 |
| e051_lncrna_func_acc | AG | 2961 | AUROC | 0.7282 | 0.7535 | 0.7487 | 0.7596 | +0.0252 | +0.0108 |
| e051_lncrna_func_acc | Enformer | 2961 | AUROC | 0.7282 | 0.7454 | 0.7487 | 0.7552 | +0.0171 | +0.0065 |
| e051_lncrna_func_acc | Borzoi | 2961 | AUROC | 0.7282 | 0.7528 | 0.7487 | 0.7616 | +0.0245 | +0.0129 |
| e051_lncrna_func_acc | RNA-FM | 2961 | AUROC | 0.7282 | 0.7471 | 0.7487 | 0.7480 | +0.0189 | -0.0007 |
| e051_lncrna_cas13 | AG | 871 | AUROC | 0.5897 | 0.6012 | 0.5532 | 0.5648 | +0.0115 | +0.0115 |
| e051_lncrna_cas13 | Enformer | 871 | AUROC | 0.5897 | 0.5753 | 0.5532 | 0.5510 | -0.0145 | -0.0022 |
| e051_lncrna_cas13 | Borzoi | 871 | AUROC | 0.5897 | 0.5818 | 0.5532 | 0.5533 | -0.0079 | +0.0001 |
| e051_lncrna_cas13 | RNA-FM | 871 | AUROC | 0.5897 | 0.5792 | 0.5532 | 0.5533 | -0.0105 | +0.0001 |
| e071_mirna_hela | AlphaGenome_disrupt | 2944 | Spearman | 0.4201 | 0.4208 | 0.4415 | 0.4420 | +0.0008 | +0.0005 |
| e071_mirna_hela | AlphaGenome_expr | 2944 | Spearman | 0.4201 | 0.4197 | 0.4415 | 0.4407 | -0.0004 | -0.0008 |
| e071_mirna_hela | RNA-FM | 2944 | Spearman | 0.4201 | 0.4202 | 0.4415 | 0.4405 | +0.0001 | -0.0010 |
| e071_mirna_hek | AlphaGenome_disrupt | 2726 | Spearman | 0.3421 | 0.3454 | 0.3927 | 0.3962 | +0.0034 | +0.0035 |
| e071_mirna_hek | AlphaGenome_expr | 2726 | Spearman | 0.3421 | 0.3413 | 0.3927 | 0.3925 | -0.0008 | -0.0002 |
| e071_mirna_hek | RNA-FM | 2726 | Spearman | 0.3421 | 0.3634 | 0.3927 | 0.3927 | +0.0213 | +0.000044 |

### Table S17c. Selected alpha by task and baseline.

| Task | Baseline | Outer folds | Alpha: fold count | At upper 1e6 |
|---|---|---|---|---|
| e051_lncrna_func | N | 24 | 10.0: 24 | 0 |
| e051_lncrna_func | NS | 24 | 100.0: 24 | 0 |
| e051_lncrna_func_acc | N | 24 | 10.0: 24 | 0 |
| e051_lncrna_func_acc | NS | 24 | 100.0: 24 | 0 |
| e051_lncrna_cas13 | N | 24 | 10.0: 19; 100.0: 5 | 0 |
| e051_lncrna_cas13 | NS | 24 | 1000.0: 24 | 0 |
| e071_mirna_hela | N | 23 | 10.0: 23 | 0 |
| e071_mirna_hela | NS | 23 | 100.0: 23 | 0 |
| e071_mirna_hek | N | 23 | 1.0: 12; 10.0: 11 | 0 |
| e071_mirna_hek | NS | 23 | 100.0: 7; 1000.0: 16 | 0 |

The grid was 0.1, 1, 10, 100, 1000, 10000, 100000 and 1000000. All composition-baseline folds selected 100 or 1000. Selected values were below the upper bound, resolving the tested grid-edge concern. Baseline optimality still depends on the candidate specification. Cas13 and HEK293FT RNA-FM increments after composition became small positive values, showing that their R7 negative signs were sensitive to regularization. See analysis/regularization for saved predictions, per-fold descriptive metrics and protocol records.

**Table S18. RiNALMo versus RNA-FM model-swap sensitivity on the same members. Both models use the same native-scale estimator and fixed-OOF evaluation boundary. This comparison extends the evaluated model set outside the primary family. For miRNA, one-way envelopes and crossed-pigeonhole ranges show sensitivity to alternative dependence constructions. The results concern these named models and remain descriptive rather than calibrated joint confirmation or a model-class estimate.**

| Member | n | Native effect | RiNALMo delta and inference | RNA-FM delta and inference |
|---|---|---|---|---|
| e051_lncrna_cas13 | 871 | delta_auroc | -0.0102; descriptive fixed-OOF locus 95% range -0.0308 to +0.0077; historical fixed-prediction tail area=0.617 | -0.0120; descriptive fixed-OOF locus 95% range -0.0235 to -0.0006; historical fixed-prediction tail area=0.915 |
| e051_lncrna_func | 2961 | delta_auroc | +0.0470; descriptive fixed-OOF locus 95% range +0.0183 to +0.0785; historical fixed-prediction tail area=<0.001 | +0.0452; descriptive fixed-OOF locus 95% range +0.0137 to +0.0780; historical fixed-prediction tail area=<0.001 |
| e051_lncrna_func_acc | 2961 | delta_auroc | +0.0174; descriptive fixed-OOF locus 95% range +0.0040 to +0.0314; historical fixed-prediction tail area=<0.001 | +0.0184; descriptive fixed-OOF locus 95% range +0.0040 to +0.0328; historical fixed-prediction tail area=0.006 |
| e071_mirna_hek | 2726 | delta_cv_spearman | +0.0238; non-joint one-way 95% resampling envelope +0.0123 to +0.0342; one-way Pmax sensitivity=<0.001; crossed confirmatory P/q not assessed; crossed-pigeonhole 95% sensitivity range +0.0059 to +0.0443; not a confirmatory interval | +0.0214; non-joint one-way 95% resampling envelope +0.0107 to +0.0306; one-way Pmax sensitivity=<0.001; crossed confirmatory P/q not assessed; crossed-pigeonhole 95% sensitivity range +0.0058 to +0.0376; not a confirmatory interval |
| e071_mirna_hela | 2944 | delta_cv_spearman | +0.0037; non-joint one-way 95% resampling envelope +0.0002 to +0.0075; one-way Pmax sensitivity=<0.001; crossed confirmatory P/q not assessed; crossed-pigeonhole 95% sensitivity range -0.0021 to +0.0095; not a confirmatory interval | +0.0002; non-joint one-way 95% resampling envelope -0.0012 to +0.0015; one-way Pmax sensitivity=0.303; crossed confirmatory P/q not assessed; crossed-pigeonhole 95% sensitivity range -0.0020 to +0.0022; not a confirmatory interval |

**Table S19. Classification matched-null stratum-resolution sensitivity. Nuisance-quantile stratum count is a parameter of the locus-permutation classification comparator. Ranges and historical tail areas describe its behavior conditional on fixed OOF predictions, without calibrated confirmatory interpretation.**

| Member | k | n | Native delta AUROC | Descriptive fixed-OOF locus range | Historical tail area | Realized blocks (min-max n) |
|---|---|---|---|---|---|---|
| e051_lncrna_func | 5 | 2961 | +0.1125 | +0.0680 to +0.1537 | <0.001 | 116 (1-58) |
| e051_lncrna_func | 10 | 2961 | +0.1125 | +0.0680 to +0.1537 | <0.001 | 231 (1-34) |
| e051_lncrna_func_acc | 5 | 2961 | +0.0252 | +0.0094 to +0.0412 | <0.001 | 116 (1-73) |
| e051_lncrna_func_acc | 10 | 2961 | +0.0252 | +0.0094 to +0.0412 | <0.001 | 231 (1-47) |

**Table S20. Record of the retired interval-derived margin and equivalence analysis. The grid was not computed because fixed-prediction descriptive ranges do not provide the inferential basis for TOST, candidate-margin containment, minimum detectable effects, power, detectability or negligibility. Small-or-absent-effect assessment therefore remains open. Provenance retains the 18-row retirement receipt to distinguish this unperformed analysis from the reported results.**

### Supplementary figures

**Figure S1. TargetScan aggregate-identity sensitivity.**

Points show all fixed-alpha baseline and augmented configurations for the exact-identity HeLa 1227 and HEK293FT 645 sets and, separately, the 705-row HEK293FT terminal-U mapping assumption. Family and mature-miRNA aggregates sum local source rows at gene level, without experimental isoform/site alignment. Baseline content and model increments are compared on identical members within each set. Within-set comparisons reveal the contribution of aggregate identity; between-set differences additionally reflect selection. Values are descriptive pooled OOF Spearman estimates without confidence intervals.

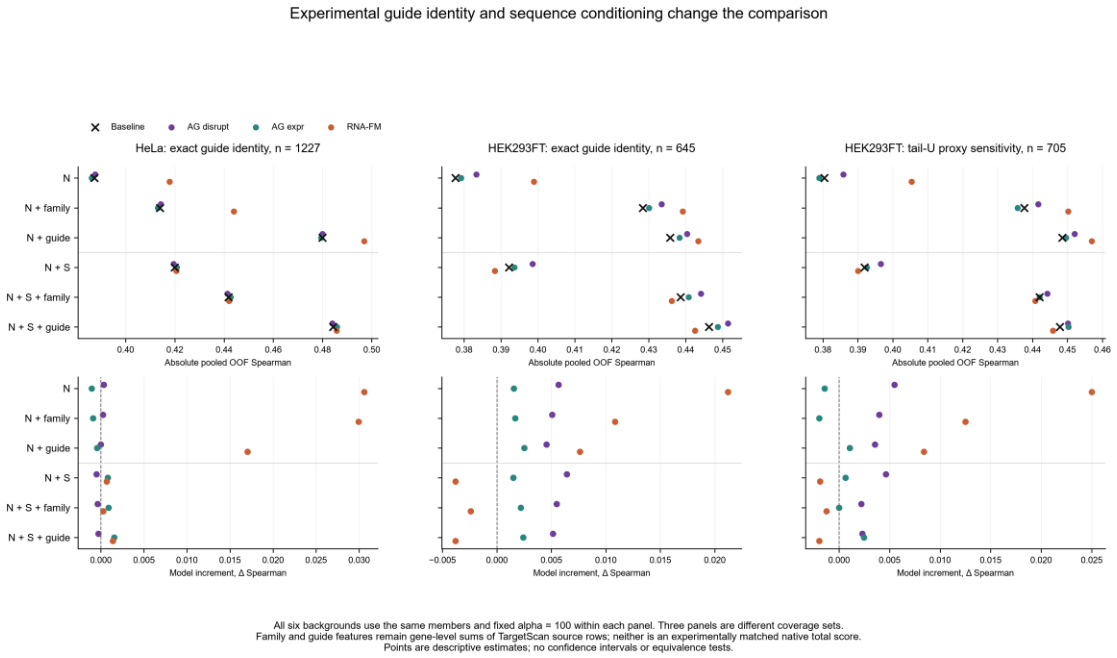


*TargetScan aggregate-identity sensitivity*

**Figure S2. Expanded regularization sensitivity.**

The R8 expanded-grid comparison retains all five tasks and 18 model cells and compares composition-conditioned increments with the initial R7 restricted-grid results. Candidate alpha extends through 1,000,000, while selected composition-baseline values are 100 or 1000. Effects remain task-native AUROC or Spearman; no inferential interval is shown. The Cas13 and HEK293FT RNA-FM sign reversals identify estimates that are sensitive to the regularization grid. Equivalence and absent-information claims remain outside this descriptive comparison.

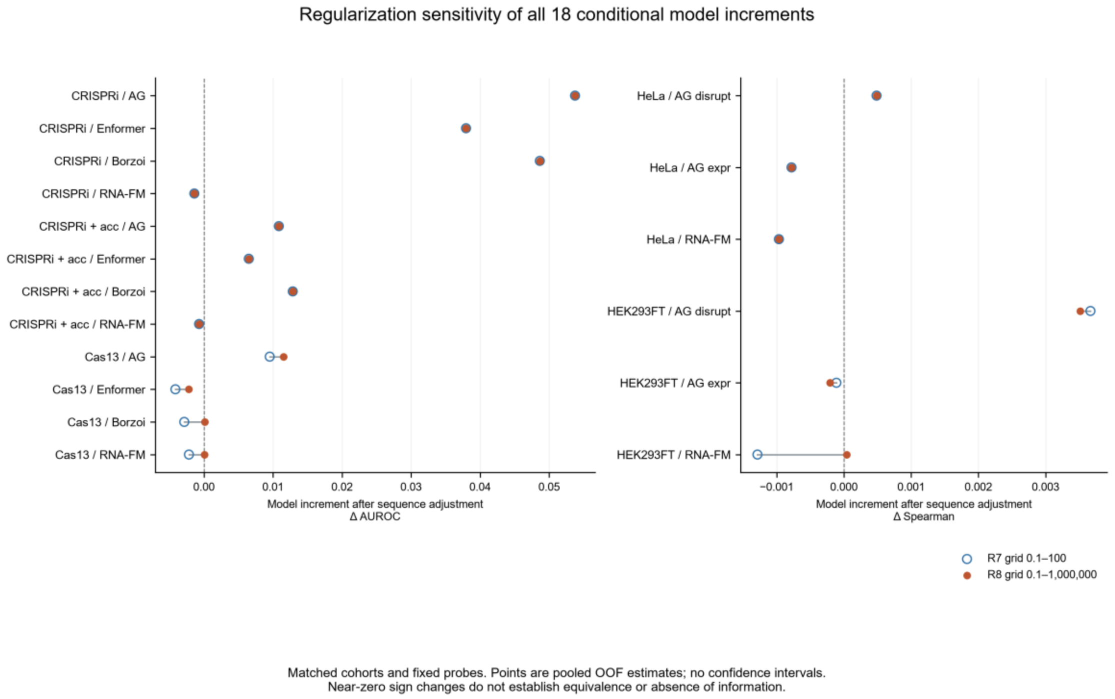


*Expanded regularization sensitivity*

### AI assistance use log

OpenAI Codex assisted with evidence-based corrections of the existing author manuscript, language editing, numerical synchronization, code execution and document preparation. The separate ledger records source text, revised text and the evidence supporting each substantive change. The authors retain responsibility for the interpretation, citations and final submission decisions.

### Additional source and assistance record

The returned evidence package adds 15 native Claude Code JSONL files (12 main sessions and three linked subagent sessions, September 2026) and 55 native Codex rollouts spanning June–August 2026. These sources cover distinct portions of the interaction history. The 391 keyword-filtered user-prompt records provide a subset of prompt history; the June Claude archive and some assistant or tool turns remain incomplete. Native September records document user authorization and subsequent producer calls. Compaction, unmatched tool calls and incomplete external-output preservation leave gaps in historical interaction and stdout recovery. The available early-session and cleanup-state records are insufficient to determine the cause of those gaps.

All 8631 current AlphaGenome summary records matched the historical evidence union; matching raw arrays still cover only 75 current requests (24 HeLa and 51 HEK293FT), with no newly recovered current raw request. For 68, the two scorers' separate original observation axes were not retained. The records thus support summary-level concordance, with raw-response reconstruction remaining partial.

The retained 2025 AlphaGenome Output Terms concern training variant-effect models similar to AlphaGenome and do not establish a blanket prohibition on every supervised probe. The accepted API Additional Terms, acceptance records and account-to-batch linkage were not supplied. Local user authorization and provider permission address separate questions. The applicable permission requirements and compliance status remain unresolved from the available documents. R8 OpenAI Codex assistance comprised local corrections of existing author text, numerical synchronization, bounded analyses and change-ledger preparation. R9 assistance added evidence review and scoped provenance amendments. The authors retain responsibility for the final interpretation and submission.
